# PPP6C negatively regulates oncogenic ERK signaling through dephosphorylation of MEK

**DOI:** 10.1101/2020.09.18.303859

**Authors:** Eunice Cho, Hua Jane Lou, Leena Kuruvilla, David A. Calderwood, Benjamin E. Turk

**Affiliations:** Department of Pharmacology, Yale School of Medicine, New Haven, CT

## Abstract

Flux through the RAF-MEK-ERK protein kinase cascade is shaped by phosphatases acting on the core components of the pathway. Despite being an established drug target and a hub for crosstalk regulation, little is known about dephosphorylation of MEK, the central kinase within the cascade. Here, we identify PPP6C, a phosphatase frequently mutated or downregulated in melanoma, as a major MEK phosphatase in cells exhibiting oncogenic ERK pathway activation. Recruitment of MEK to PPP6C occurs through an interaction with its associated regulatory subunits. Loss of PPP6C causes hyperphosphorylation of MEK at both activating and crosstalk phosphorylation sites, promoting signaling through the ERK pathway and resistance to the growth inhibitory effects of MEK inhibitors. Recurrent melanoma-associated PPP6C mutations cause MEK hyperphosphorylation, suggesting that they promote disease at least in part by activating the core oncogenic pathway driving melanoma. Collectively, our studies identify a key negative regulator of ERK signaling that may influence susceptibility to targeted cancer therapies.

## INTRODUCTION

The RAF-MEK-ERK mitogen-activated protein kinase (MAPK) signaling cascade regulates essential cellular processes including cell proliferation, differentiation, and survival(Lavoie et al., 2020). Deregulation of ERK signaling, typically through mutations in core pathway components or upstream regulators, is among the most frequent driver events in human cancer (Burotto et al., 2014). Malignant melanoma in particular is dependent upon hyperactivate ERK signaling. About half of human melanomas harbor mutations in the BRAF gene, the most common being the V600E mutation that causes high level constitutive activity (Cancer Genome Atlas Network, 2015; Davies et al., 2002; Hayward et al., 2017; Hodis et al., 2012; Krauthammer et al., 2015). Most other melanoma tumors have either gain-of-function mutations in the NRAS GTPase, a direct activator of RAF kinases, or loss-of-function mutations in the RAS GTPase activating protein NF1. The dependence of melanomas on the ERK pathway fueled the development and approval of selective BRAF inhibitors (BRAFi) and MEK inhibitors (MEKi) with clinical efficacy in treating tumors harboring BRAF^V600E^ mutations (Samatar and Poulikakos, 2014). Unfortunately, responses to BRAFi and MEKi are almost invariably short-lived due to the development of acquired resistance (Flaherty et al., 2012). While resistance to these agents can involve activation of alternative cell growth and survival pathways, it most commonly occurs through reactivation of the ERK pathway despite the continued presence of inhibitor(Lim et al., 2017). Mechanisms underlying ERK pathway reactivation include acquisition of RAS or MEK mutations, BRAF amplification or alternative splicing, disruption of negative feedback regulation, and induction of receptor tyrosine kinases (Johnson et al., 2015; Rizos et al., 2014; H. Shi et al., 2014; Van Allen et al., 2014). Understanding how tumor cells become resistant to BRAFi and MEKi can suggest additional therapeutic targets and thus contribute to the development of durable and more generally applicable cancer treatments. In addition, investigations into mechanisms of inhibitor resistance have provided insight into the basic wiring of the ERK pathway and how it participates in larger signaling networks.

Because proper control of the ERK pathway is important to normal physiology, the core cascade is positioned within a complex network involving extensive feedback and crosstalk regulation (Lavoie et al., 2020). By counteracting regulatory phosphorylation events, protein phosphatases play key roles in controlling the magnitude and duration of ERK signaling, and their dysregulation can contribute to disease and influence inhibitor sensitivity. For example, ERK signaling induces expression of dual-specificity MAPK phosphatases (DUSPs), which dephosphorylate and inactivate ERK (Bermudez et al., 2010). Disruption of this negative feedback loop through deletion or downregulation of ERK-selective DUSPs has been reported in some tumors and is associated with more advanced disease and poor patient prognosis (Cai et al., 2015; Okudela et al., 2009; Shin et al., 2013; S. Xu et al., 2005). Protein phosphatases also have important roles in positively regulating ERK signaling. For example, the MRAS-SHOC2-PP1 complex dephosphorylates an inhibitory site on RAF and mediates ERK pathway reactivation induced by MEKi in KRAS mutant pancreatic and lung cancers (Jones et al., 2019; Sulahian et al., 2019). Germline gain-of-function mutations in MRAS and SHOC2 are a cause of developmental disorders termed RASopathies that are characterized by hyperactive ERK signaling (Young et al., 2018). Protein phosphatase 2A (PP2A) can promote ERK signaling by dephosphorylating inhibitory feedback phosphorylation sites on RAF and the ERK pathway scaffold KSR1 (Ory et al., 2003). While these phosphatases regulating RAF and ERK have established roles in normal and pathological signaling, less is known about phosphatases regulating MEK, the central component of the cascade. PP2A was initially identified as a MEK phosphatase *in vitro* and in non-transformed monkey kidney CV-1 cells (Gómez and Cohen, 1991; Sontag et al., 1993). However, other studies have suggested that PP2A restrains oncogenic MAPK signaling primarily through direct dephosphorylation of ERK.

Here we have identified PPP6C as a MEK phosphatase that negatively regulates oncogenic ERK signaling and promotes sensitivity to MEKi. The potential significance of this observation is underscored by the presence of loss-of-function PPP6C mutations in 6-9% of malignant melanomas, generally co-occurring with BRAF and NRAS mutations (Cancer Genome Atlas Network, 2015; Hodis et al., 2012; Krauthammer et al., 2015). PPP6C is a conserved essential serine-threonine metallophosphatase related to the catalytic subunit of PP2A (Y. Shi, 2009). PPP6C functions within heterotrimeric PP6 holoenzymes consisting of a Sit4-associated protein (SAPS) domain regulatory subunit (PPP6R1, PPP6R2 or PPP6R3) that mediates substrate recruitment and an ankyrin repeat (ANKRD28, ANKRD44 or ANKRD52) subunit that may serve a scaffolding role (Stefansson and Brautigan, 2006; Stefansson et al., 2008). The presence of multiple regulatory and scaffolding subunits defines nine potential PP6 complexes that collectively participate in a variety of cellular processes, including cell cycle progression (Douglas et al., 2014; Rusin et al., 2017; Zeng et al., 2010), the DNA damage response (Douglas et al., 2014; Hosing et al., 2012; Shen et al., 2011; Zhong et al., 2011), autophagy (Wengrod et al., 2015), miRNA processing (Golden et al., 2017), inflammatory response (Kajino et al., 2006; Ye et al., 2015), and antiviral immunity (Tan et al., 2017). Relevant to its role as a tumor suppressor, loss of PPP6C causes defects in mitotic spindle assembly owing to hyperphosphorylation of the kinase Aurora A, and consequent genomic instability has been proposed as an early event in tumorigenesis (Gold et al., 2014; Hammond et al., 2013). By contrast, connections between PPP6C and core oncogenic pathways have yet to be described.

We identified PPP6C as a MEK phosphatase through an shRNA screen to identify genes modulating the response of a BRAF mutant melanoma cell line to MEKi. Loss of PPP6C appears to cause MEKi resistance by inducing MEK hyperphosphorylation, thereby maintaining ERK activity in the presence of inhibitor. We found that negative regulation of MEK by PPP6C appears to be a general feature of cells characterized by hyperactive ERK signaling, including cell lines harboring NRAS and KRAS mutations. Furthermore, we find that melanoma-associated mutations impair the ability of PPP6C to dephosphorylate MEK and promote resistance to MEKi. These results provide a likely explanation for the prevalence of PPP6C mutations in melanomas and potentially other tumors, and suggest a potential mechanism for inhibitor resistance. Furthermore, these studies report a major negative regulator that dephosphorylates the central component in the core ERK signaling cascade.

## RESULTS

### Pooled shRNA library screen identifies PPP6C as a mediator of response to MEKi

To identify genes involved in modulating the response to MEK inhibitors, we performed a pooled shRNA screen in the BRAF^V600E^ mutant melanoma cell line 501mel (Figure 1A). We used a custom-made lentiviral library of 7,649 shRNAs targeting 817 genes encoding annotated protein and lipid kinases and phosphatases (Table S1). After transduction with the shRNA library, we harvested a portion of the cells for a start time reference sample (T_0_) and divided the remaining cells into 5 populations, which were subsequently treated with either vehicle control or one of two concentrations of the MEKis trametinib (1 nM and 3.3 nM) or selumetinib (33 nM and 100 nM). MEKi concentrations were chosen to flank the approximate IC_50_ for inhibition of cell growth. Cells were propagated through 10 population doublings (T_1_-T_10_). The change in abundance of each shRNA under each growth condition was then determined by next-generation sequencing (Illumina HiSeq) following PCR amplification from genomic DNA preparations of cells collected at T_0_ and T_10_ (Table S1). As expected for a cell line carrying a BRAF^V600E^ mutation, hairpins targeting BRAF were depleted in all populations over time (Figure 1C), providing an internal control for the screens.

**Figure 1.**
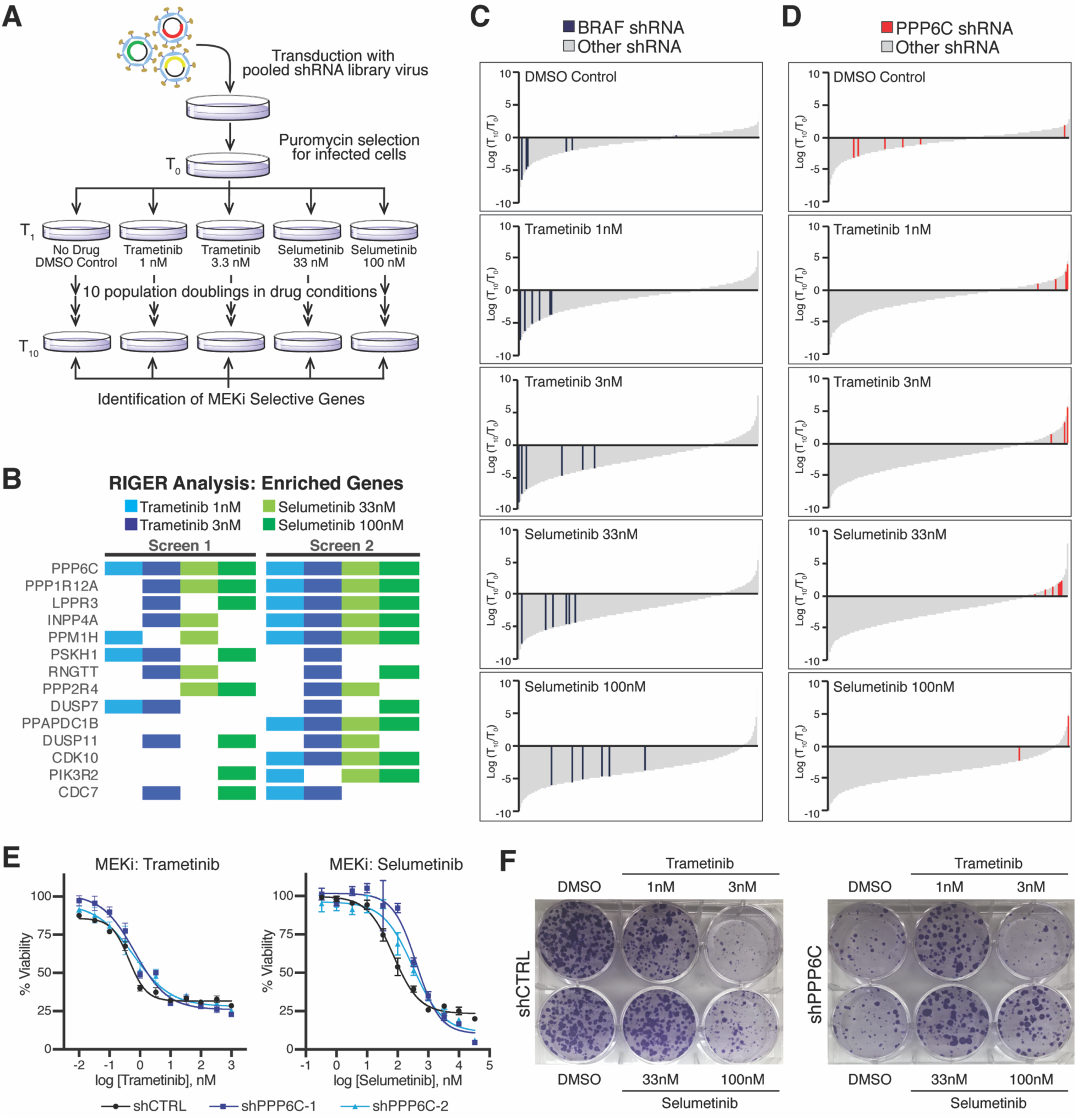
Pooled shRNA library screen identifies PPP6C as a mediator of response to MEKi. (A) Schematic of the pooled shRNA MEKi sensitivity screen. (B) Top enriched genes for each drug condition from two replicates of the screen. Colored boxes indicate the genes ranked in the top 50 (top 6.1%) enriched genes by RIGER for that drug condition not found in the DMSO control condition. (C) Changes in all shRNA hairpins shown as log_2_(T_10_/T_0_) from most depleted to most enriched for each drug condition. Bars representing shRNA hairpins targeting BRAF are shown in blue. All others are shown in grey. (D) Changes in all shRNA hairpins shown as log_2_(T_10_/T_0_) arranged as in (C), but with red bars indicating shRNA hairpins targeting PPP6C. (E) 501mel cell lines stably expressing control shRNA (shCTRL) or PPP6C-targeting shRNA (shPPP6C-1, shPPP6C-2) were treated for 72 hours with increasing concentrations of trametinib or selumetinib. Cell viability was detected by alamarBlue reagent and normalized to a no drug control for each cell line. Dose response curves for shCTRL (black), shPPP6C-1 (dark blue), and shPPP6C-2 (light blue) are shown. (F) shCTRL, shPPP6C-1, and shPPP6C-2 expressing 501mel cells were cultured in DMSO or the indicated concentration of trametinib or selumetinib for 2 weeks in colony forming assays. Colonies were stained with crystal violet.

We used RNAi gene enrichment ranking (RIGER) analysis (Luo et al., 2008) to rank genes based on depletion or enrichment of their shRNAs in the screen. This analysis revealed that hairpins targeting PPP6C were the most consistently enriched in the presence of MEKi (Figures 1B and S1). PPP6C was the top ranked gene in 5 of 8 MEKi conditions across the two screens and scored within the top 20 genes under all drug conditions (Table S2). While PPP6C-targeting hairpins highly enriched in the presence of MEKi, they were generally depleted from the untreated culture (Figure 1D). These results suggest that PPP6C is required both for optimal cell growth and for a maximal cytostatic response to MEKi.

To confirm a role for PPP6C in conferring resistance to MEKi, 501mel cells were transduced with non-targeting control shRNA (shCTRL) or one of two independent shRNAs (shPPP6C-1, shPPP6C-2) that efficiently decrease PPP6C expression (see Figure 2A). PPP6C knockdown caused modest but consistent rightward shifts to MEKi dose responses curves, increasing IC_50_ values for growth inhibition by trametinib and selumetinib (Figure 1E). In clonogenic assays, cells transduced with control shRNA exhibited dose-dependent growth inhibition by both MEKi as anticipated (Figure 1F). PPP6C knockdown alone substantially decreased cell growth, yet this growth defect was largely reversed in the presence of low concentrations of MEKi. In addition, cells in which PPP6C had been silenced grew slightly better than control cells at higher MEKi concentrations. These observations verify that PPP6C silencing renders 501mel cells less sensitive to the cytostatic effects of MEKi, consistent with the behavior of shRNAs targeting PPP6C in the screens. The enhanced growth observed upon MEKi treatment of PPP6C knockdown cells is consistent with ERK pathway inhibitor addiction, a phenomenon observed in other settings of MEKi and BRAFi resistance (Hong et al., 2018; Kong et al., 2017; Moriceau et al., 2015).

**Figure 2.**
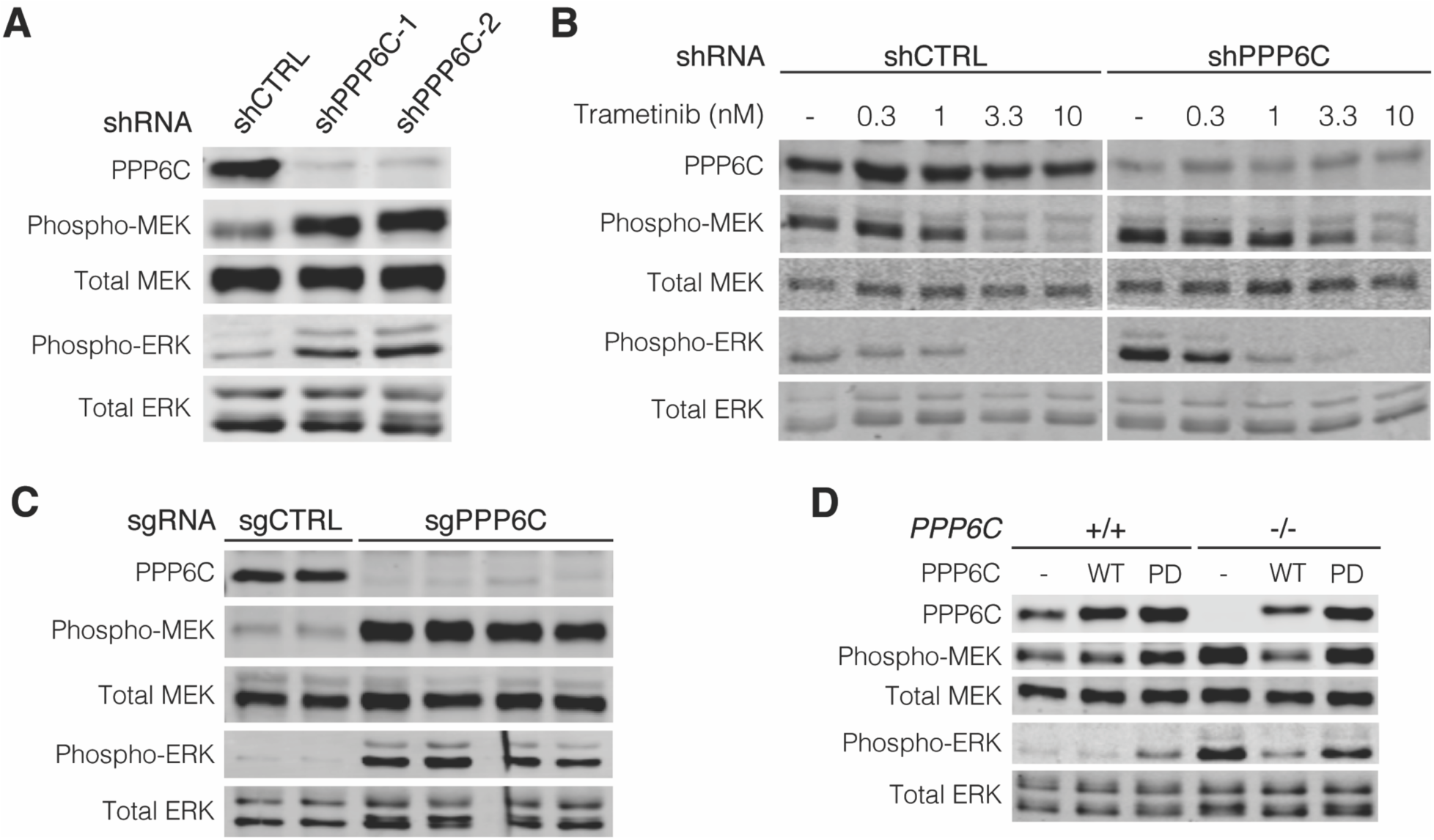
PPP6C negatively regulates ERK signaling. (A) 501mel cells stably expressing shCTRL, shPPP6C-1, and shPPP6C-2 were lysed and assessed by immunoblot for levels of total and phosphorylated MEK and ERK. (B) shCTRL and shPPP6C expressing 501mel cells were treated with the indicated concentrations of trametinib for 1 hour and lysed. Phosphorylated and total levels of MEK and ERK were detected by immunoblot. (C) *PPP6C*^*-/-*^ 501mel cell lines were established by CRISPR/Cas9. *PPP6C*^*+/+*^ and *PPP6C*^*-/-*^ cell lines were lysed and assessed by immunoblot for phospho and total levels of MEK and ERK. (D) *PPP6C*^*+/+*^ and *PPP6C*^*-/-*^ 501mel cells stably expressing WT PPP6C, phosphatase inactive PPP6C^D84N^ (PD), or GFP (-) as a control were lysed and assessed by immunoblot for phosphorylated and total MEK and ERK.

### PPP6C negatively regulates ERK signaling

Resistance to ERK pathway inhibitors can occur through multiple mechanisms, for example through activation of bypass pathways (Johnson et al., 2015; Lu et al., 2017; H. Shi et al., 2014). Indeed, in the shRNA screen, hairpins targeting two negative regulators of phosphoinositide 3-kinase (PI3K) signaling, *PIK3R2* and *INPP4A* (Table S1), were enriched in MEKi-treated cells, suggesting that activation of this pathway can cause MEKi resistance in 501mel cells. We suspected that loss of PPP6C led to ERK pathway hyperactivation, as this phenomenon can underlie both inhibitor resistance and addiction (Hong et al., 2018; Moriceau et al., 2015). Indeed, we observed that silencing of PPP6C in 501mel cells elevated the levels of activating phosphorylation of ERK1/2 and MEK1/2 (Figure 2A). Cells expressing shPPP6C required higher concentrations of inhibitor to reduce MEK and ERK phosphorylation to levels seen in control cells (Figure 2B), likely explaining why PPP6C knockdown causes low level MEKi resistance. Likewise, low concentrations of MEKi attenuate hyperactivation of ERK signaling that is presumably toxic to these cells, explaining why reduced PPP6C expression leads to inhibitor addiction. Furthermore, the ability of low concentrations of MEKi to rescue growth (Figure 1F) suggests that PPP6C is required for optimal growth of 501mel cells largely because it restrains ERK signaling.

To further verify that PPP6C regulates ERK signaling, we generated clonal PPP6C knockout 501mel lines by CRISPR/Cas9-mediated gene disruption (Figures 2C and S2A). Notably, we were unable to expand *PPP6C*^-/-^ clones unless we supplemented the growth media with a low concentration of the MEKi trametinib, and as with PPP6C knockdown these cells displayed inhibitor addiction (Figure S2B). The complete loss of PPP6C in these clones resulted in an even more pronounced increase in MEK and ERK phosphorylation than seen with partial loss of PPP6C via shRNA (Figure 2C). Re-expression of wild-type (WT) PPP6C, but not a phosphatase inactive mutant (D84N, PD) in these cells reversed ERK hyperactivation (Figure 2D), indicating that negative regulation of ERK signaling requires PPP6C phosphatase activity.

To determine whether PPP6C acts as a general regulator of ERK signaling outside of the cell line used for our screen, we examined the effect of PPP6C knockdown in a panel of cell lines of varying genotype and lineage (Figure 3A). We found that silencing PPP6C expression led to MEK hyperphosphorylation in each of five additional *BRAF*^*V600*^ mutant melanoma cell lines tested, including YURIF cells harboring a *PPP6C*^*S270L*^ mutation. Among four *NRAS*^*Q61*^ mutant melanoma cell lines, all but one (YUGASP) exhibited increased MEK and ERK phosphorylation upon PPP6C knockdown. We observed the same phenomenon with MEL-ST, a non-transformed immortalized melanocyte cell line. Additionally, in cell lines derived from other tumor types with *BRAF* and *RAS* mutations, including a lung adenocarcinoma cell line and three colon carcinoma cell lines, we also observed increased MEK and ERK phosphorylation with PPP6C loss. The osteosarcoma cell line U2OS, which does not harbor *BRAF* or *RAS* mutations, was unaffected by PPP6C knockdown. Overall, the large majority of cell lines examined displayed PPP6C regulation of ERK signaling.

**Figure 3.**
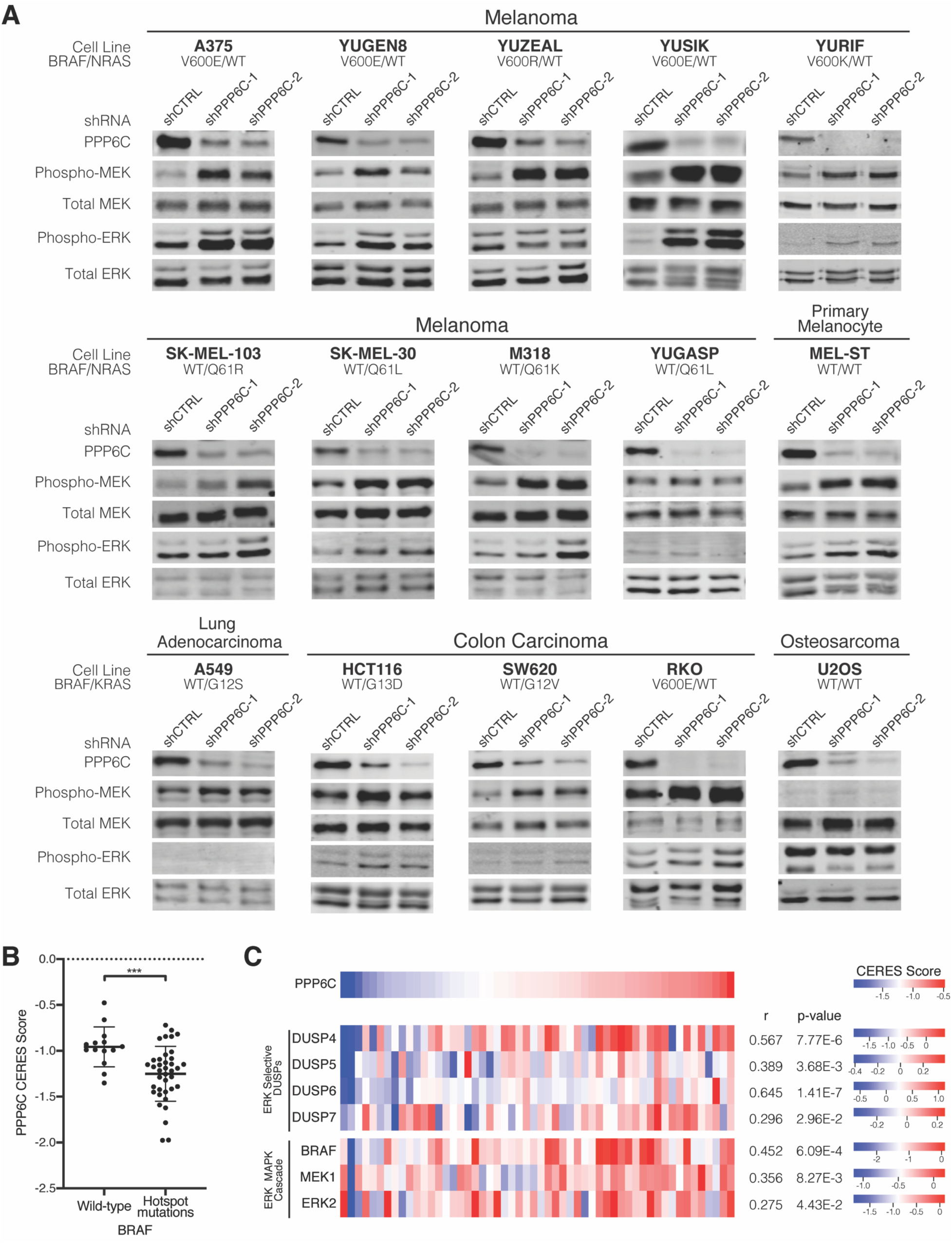
PPP6C regulation of ERK signaling is prominent in ERK pathway-driven cancer cells. (A) The indicated cell lines were transduced to stably express shCTRL, shPPP6C-1, or PPP6C-2. Cells were lysed and phosphorylated and total levels of MEK and ERK were analyzed by immunoblotting. (B) PPP6C CERES scores for skin cancer cell lines with WT BRAF or hotspot BRAF mutations from the Cancer Dependency Map Project. Cell lines harboring non-recurrent BRAF variants of unknown significance were excluded. Data are represented as mean ± SD. \*\*\**p* < 0.0005, Welch’s t test. (C) Heatmaps depicting CERES scores of PPP6C, ERK selective DUSPs, and ERK MAPK cascade components in skin cancer cell lines using data from Cancer Dependency Map Project. Pearson’s correlation coefficients and *p*-values from linear regression analysis of each gene with PPP6C in the DepMap portal are listed.

We further examined a role for PPP6C as a regulator of ERK signaling by analyzing data from the Cancer Dependency Map Project (Dempster et al., 2019; DepMap, 2020; Meyers et al., 2017), which compiles the results of genome-wide CRISPR/Cas9 screens across a large panel (739) of cell lines. *PPP6C* is categorized as a common essential gene with an average CERES gene dependency score of −0.10 ± 0.28, where a more negative score indicates a larger effect on cell growth or survival (Meyers et al., 2017). Notably, in skin cancer cell lines, the mean CERES score for PPP6C is −1.16 ± 0.31, indicating these cell lines are in general more dependent on *PPP6C*. Among skin cancer cell lines, those with BRAF hotspot mutations were significantly more dependent on PPP6C than those that are wild-type for BRAF (Figure 3B). Taken together with our data in 501mel cells, these data suggest that cells characterized by hyperactive ERK signaling are more sensitive to loss of PPP6C.

We further compared the dependency of skin cancer cell lines on PPP6C and core components of the ERK signaling pathway by examining pairwise correlations between genes. In these data, co-dependency between two genes across cell lines can indicate that they participate in a common pathway(Boyle et al., 2018; Kim et al., 2019). Dependency on *PPP6C* significantly correlates with dependency on *BRAF, MAP2K1* (encoding MEK1), and *MAPK1* (encoding ERK2) (Figures 3C and S3A-D). Strikingly, strongest co-dependency with *PPP6C* was found among negative regulators of the pathway, the ERK selective dual-specificity protein phosphatases (*DUSP4, DUSP5, DUSP6*, and *DUSP7*). Consistent with a key role for PPP6C in dephosphorylating Aurora A (Hammond et al., 2013; Zeng et al., 2010), *PPP6C* dependency negatively correlated with that of *AURKA* among the full set of cell lines across all lineages (Pearson’s correlation coefficient = −0.216, *p* = 1.40 × 10^−9^) (Figure S3F). However, this correlation was not significant in skin cancer cell lines (*p* = 0.44) (Figure S3E), suggesting that regulation of Aurora A is not the primary determinant of PPP6C dependency in these cells. Overall, these correlations suggest that in skin cancer cell lines, dependency on PPP6C is associated with its role as a regulator of ERK signaling.

### PPP6C regulates ERK signaling via MEK1/2

Hyperphosphorylation of MEK observed with PPP6C loss suggests PPP6C regulates ERK signaling either at the level of MEK or upstream of MEK. To determine which component of the ERK signaling cascade is regulated by PPP6C, we initially investigated the RAF kinases directly upstream of MEK. In BRAF^V600E^ mutant melanoma, oncogenic signaling is driven primarily by mutant BRAF, with little contribution from the other RAF isoforms ARAF and CRAF (Karasarides et al., 2004). However, in settings of BRAFi/MEKi resistance, ERK pathway reactivation can occur in a manner dependent on CRAF, for example by induction of upstream receptor tyrosine kinases (Montagut et al., 2008; H. Shi et al., 2011; Villanueva et al., 2010). To determine if increased MEK phosphorylation observed with PPP6C loss is due to compensation by ARAF or CRAF, we silenced each of the RAF isoforms by siRNA in combination with shRNA knockdown of PPP6C (Figure 4A). In both the shCTRL and shPPP6C cells, only BRAF knockdown decreased MEK phosphorylation, while silencing ARAF and CRAF alone or in combination (Figure S4) did not. Thus, in the context of PPP6C loss, BRAF remains the principal activator of MEK.

**Figure 4.**
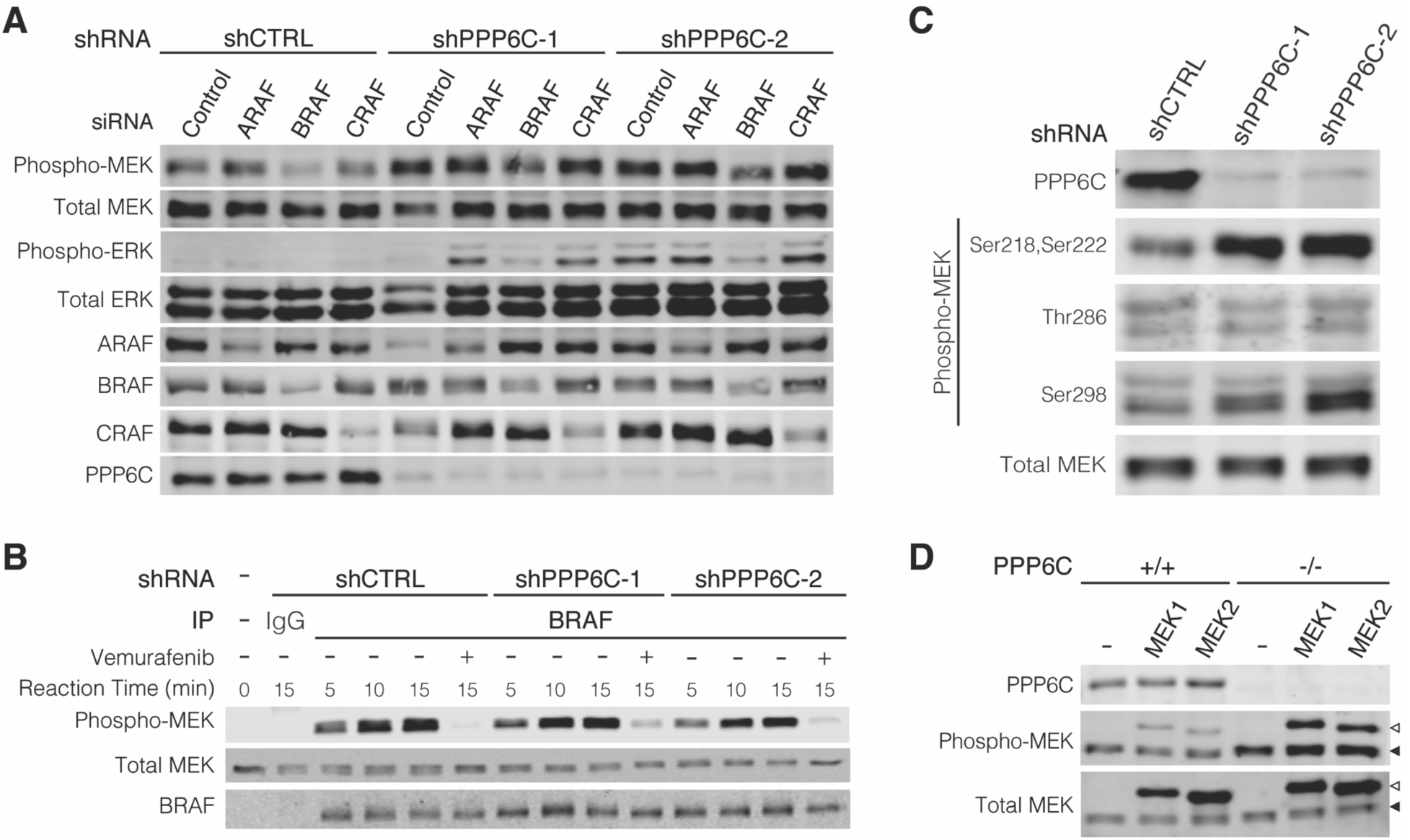
PPP6C regulates ERK signaling via MEK1/2. (A) shCTRL, shPPP6C-1, and shPPP6C-2 expressing 501mel cells were transfected with non-targeting control siRNA or siRNAs directed to ARAF, BRAF, or CRAF as indicated. Cells were lysed and assessed by immunoblot for phosphorylated and total MEK and ERK. (B) BRAF was immunoprecipitated from 501mel cells expressing shCTRL, shPPP6C-1, or shPPP6C and evaluated *in vitro* in kinase assays on MEK1 over the indicated time course. Vemurafenib (1 μM) was added to negative control reactions. Reactions were evaluated by immunoblot. (C) 501mel cells expressing shCTRL, shPPP6C-1, and shPPP6C-2 were lysed and assessed by immunoblot for MEK phosphorylation at Ser218/Ser222, Thr286, and Ser298. Non-specific cross-reacting bands in the pThr286 and pSer298 blots are indicated with an asterisk. (D) *PPP6C*^*+/+*^ and *PPP6C*^*-/-*^ 501mel cell lines were transiently transfected to express His epitope tagged MEK1 or MEK2. Cell lysates were analyzed by immunoblot for phosphorylated and total MEK. Upper bands (open arrows) correspond to ectopically expressed His-tagged MEK1/2, and lower bands (solid arrows) show endogenous MEK1/2.

We next considered whether loss of PPP6C leads to increased BRAF activity. To do so, we immunoprecipitated endogenous BRAF from shCTRL cells and shPPP6C cells and examined its vemurafenib-sensitive kinase activity on MEK1 *in vitro*. We found that BRAF isolated from both shCTRL and shPPP6C cells phosphorylated MEK1 at similar rates (Figure 4B), indicating PPP6C does not regulate BRAF activity. While we did not observe changes in RAF expression levels with PPP6C loss (Figures 4A and 4B), we did note upward electrophoretic mobility shifts suggestive of a change in the phosphorylation states of BRAF and CRAF that could have a regulatory role. We found however that treatment of cells with MEKi or BRAFi caused the multiple BRAF species to collapse into a lower, presumably less phosphorylated species (Figure S5). The increased RAF phosphorylation observed in shPPP6C cells is therefore presumably a consequence of increased negative feedback phosphorylation due to ERK hyperactivation (Lake et al., 2016; Ritt et al., 2010). In keeping with reports that BRAF^V600E^ is insensitive to feedback phosphorylation (Ritt et al., 2010), its hyperphosphorylation did not affect its activity *in vitro* and is thus unlikely to impact MEK phosphorylation in cells.

PPP6C regulates the level of RAF-mediated MEK activation loop phosphorylation without affecting the activity of RAF itself. MEK activation loop phosphorylation can be influenced by crosstalk regulation through phosphorylation at other sites, which could be subject to regulation by PPP6C. Indeed, we found that MEK1 phosphorylation at Ser298, which is mediated by PAK1 to promote activation loop phosphorylation (Coles and Shaw, 2002; Lake et al., 2016), was elevated in cells lacking PPP6C (Figure 4C). In contrast, there was no effect on phosphorylation at Thr286, a negative regulatory site phosphorylated by CDK1 or CDK5 (Rossomando et al., 1994; Sharma et al., 2002). Because regulation by Ser298 phosphorylation is specific to MEK1 and not MEK2, we examined whether PPP6C selectively regulates MEK isoforms. We found that ectopically expressed MEK1 and MEK2 (upper bands), like endogenous MEK1/2 (lower bands), were both hyperphosphorylated when expressed in *PPP6C*^*-/-*^ cells in comparison to WT cells (Figure 4D). PPP6C therefore does not preferentially regulate one isoform of MEK but instead regulates both MEK1 and MEK2. This suggests that PPP6C regulates MEK activity by modulating activation loop phosphorylation independently of crosstalk pathways.

### MEK1/2 is a direct substrate of PP6

As our findings above indicate PPP6C regulates MEK1/2 activation loop phosphorylation without affecting RAF activity, PPP6C likely promotes MEK1/2 dephosphorylation, possibly acting directly. To assess PPP6C dephosphorylation of MEK, we isolated PP6 complexes by affinity purification from HEK293T cells ectopically expressing FLAG epitope-tagged PPP6C with PPP6R3 and ANKRD28. Complexes containing WT PPP6C dephosphorylated the activation loop residues (pSer218 and pSer222) of MEK1 in a manner sensitive to the pan-PP2A family phosphatase inhibitor okadaic acid (Figure 5A). Phosphatase inactive PPP6C^D84N^ complexes had no activity against MEK1, indicating that our preparations were not contaminated with other MEK phosphatase activities. We found that PP6 also dephosphorylated pSer298 on MEK1, albeit with slower kinetics than with the activation loop sites, while it did not dephosphorylate pThr286. Thus, PP6 dephosphorylates MEK1 selectively at the same sites that are elevated in cells lacking PPP6C (Figure 4C). Furthermore, PP6 had no activity on phospho-ERK2, consistent with PPP6C acting as a regulator of MEK (Figure 5B). Overall, these studies demonstrate the direct dephosphorylation of MEK1 by PP6 with substrate and phosphorylation site specificity (Figure 5C).

**Figure 5.**
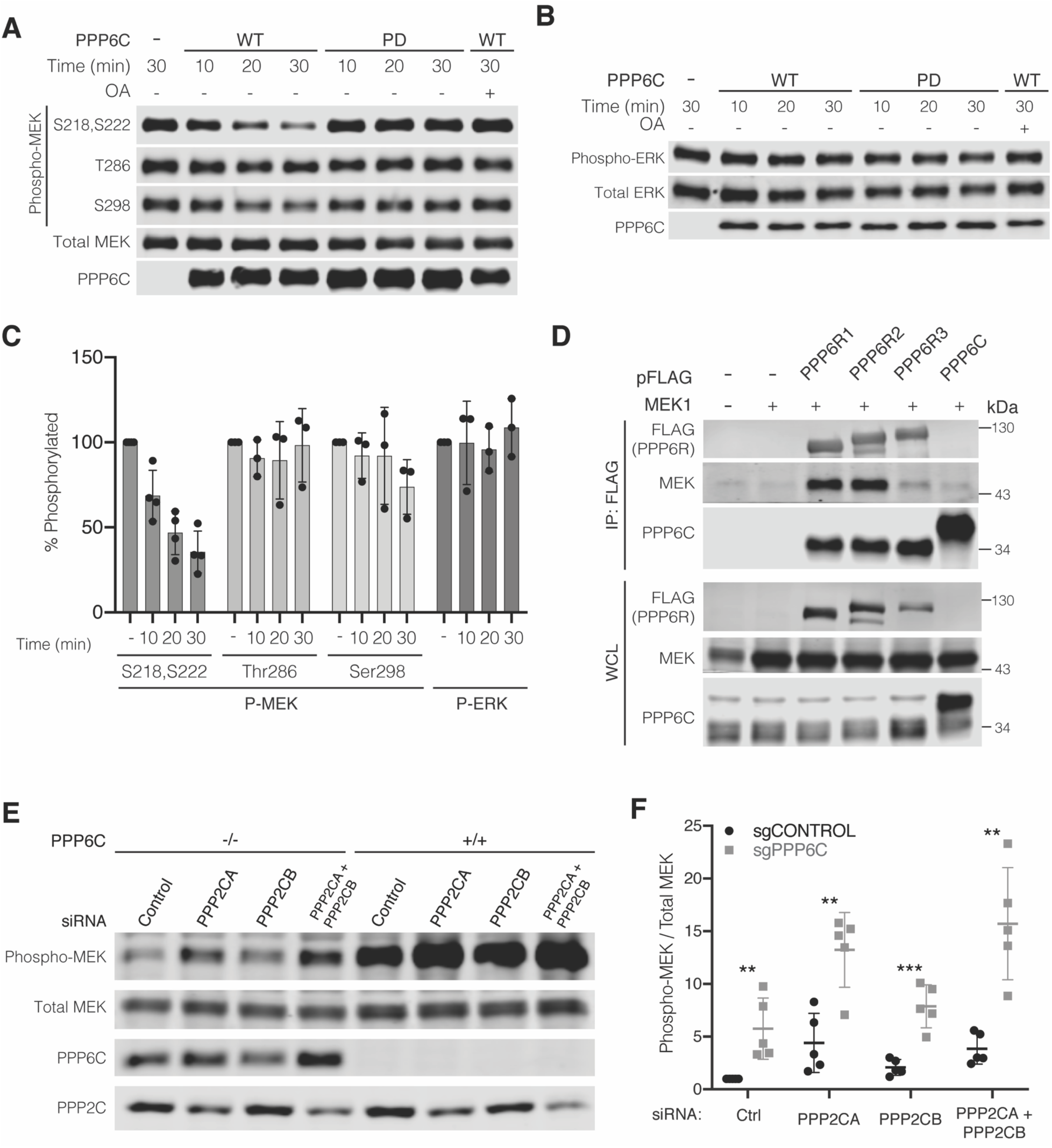
MEK1/2 is a direct substrate of PP6. (A) PP6 complexes with WT or phosphatase inactive PPP6C (PD) were partially purified from HEK293T cells and incubated with phosphorylated MEK1 *in vitro* for the indicated times. Okadaic acid (OA, 100nM) was added where indicated. Reactions were evaluated by immunoblot. (B) *In vitro* phosphatase assays evaluating phospho-ERK2 as a PP6 substrate were carried out as in (A). Reactions were evaluated by immunoblot. (C) Quantification of *in vitro* phosphatase assays in (A) and (B). Remaining phosphorylation is shown relative to the 30 min control reaction. Data are represented as mean ± SD. For MEK1 pSer218/pSer222, *n* = 4; for all other data, *n* = 3. (D) HEK293T cells were co-transfected to express the indicated FLAG epitope tagged PP6 subunit and untagged MEK1. Anti-FLAG immunoprecipitates and whole cell lysates (WCL) were evaluated by immunoblot for MEK. (E) *PPP6C*^*+/+*^ and *PPP6C*^*-/-*^ 501mel cells were transfected with non-targeting control siRNA or siRNA SMARTpools targeting PPP2CA and/or PPP2CB. Cells were lysed and evaluated by immunoblot for phosphorylated and total MEK. (F) Quantification of the relative level of MEK phosphorylation for PPP2CA/PPP2CB knockdown in *PPP6C*^*+/+*^ and *PPP6C*^*-/-*^ 501mel cells (E). MEK phosphorylation was normalized to *PPP6C*^*+/+*^, siRNA Control. Data are represented as mean ± SD, n = 5. \*\**p* < 0.01, \*\*\**p* < 0.001, unpaired t test.

To provide additional evidence that PPP6C acts directly on MEK, we performed co-immunoprecipitation experiments to determine if PP6 can interact with MEK in cells. HEK293T cells were transfected with plasmids expressing a FLAG epitope tagged PP6 subunit (PPP6C, PPP6R1, PPP6R2, or PPP6R3) and untagged MEK1. We found that MEK1 co-immunoprecipitated with each of the PP6 regulatory subunits (Figure 5D), with the amount of associated MEK1 proportional to the PPP6R expression level. Significantly less MEK1 associated with FLAG-tagged PPP6C. These results suggest that PP6 regulatory subunits serve to recruit MEK to the PP6 complex for dephosphorylation by PPP6C, consistent with a general role for non-catalytic subunits of PP6 and other PP2A family phosphatases in substrate binding (Brautigan and Shenolikar, 2018; Heo et al., 2020; Hosing et al., 2012; Stefansson and Brautigan, 2006; Zhong et al., 2011).

Our observations collectively suggest that PP6 has a general role as a MEK phosphatase across multiple cell types. Classical studies, however, had implicated PP2A as the major MEK phosphatase, suggesting that PP6 may act indirectly by regulating PP2A activity on MEK (Gómez and Cohen, 1991; Sontag et al., 1993). We therefore investigated the impact of PP6 and PP2A loss, alone and in combination, on MEK phosphorylation. We used siRNA SMARTpools to knockdown the two PP2A catalytic subunits (PPP2CA and PPP2CB) in *PPP6C*^*-/-*^ and control 501mel cells (Figure 5E). Loss of each of the two PP2A catalytic subunit isoforms individually and in combination increased MEK phosphorylation levels in the control cell line, with silencing of PPP2CA having the predominant effect. In *PPP6C*^*-/-*^ cells, PP2A downregulation further increased MEK phosphorylation, with the two phosphatases having an apparently additive effect. Notably, loss of either phosphatase increased expression levels of the other, suggestive of a compensatory mechanism (Figure 5F). This experiment suggests that PP6 and PP2A act independently to dephosphorylate MEK.

### Cancer-associated PPP6C mutations abrogate PP6 phosphatase activity against MEK1/2

PPP6C mutations are found across multiple cancer types but are most common in melanoma and other skin cancers, where they are thought to contribute to tumor development (Hodis et al., 2012; Krauthammer et al., 2015; Malicherova et al., 2019; Palmieri et al., 2018). (Hammond et al., 2013). Prior characterization of PPP6C mutations has focused primarily on non-recurrent mutations that cluster at the catalytic center, which reportedly reduce or eliminate phosphatase activity. Interestingly, there are several PPP6C hotspot residues that are recurrently mutated, with R264C being the most common (Figure 6A). When modeled onto the X-ray crystal structure of a PPP5C-peptide complex (Oberoi et al., 2016), sites of recurrent mutations are generally located within or proximal to the catalytic cleft (Figure 6B). Of these, His55 appears critical for activity as it coordinates one of the bound metal ions.

**Figure 6.**
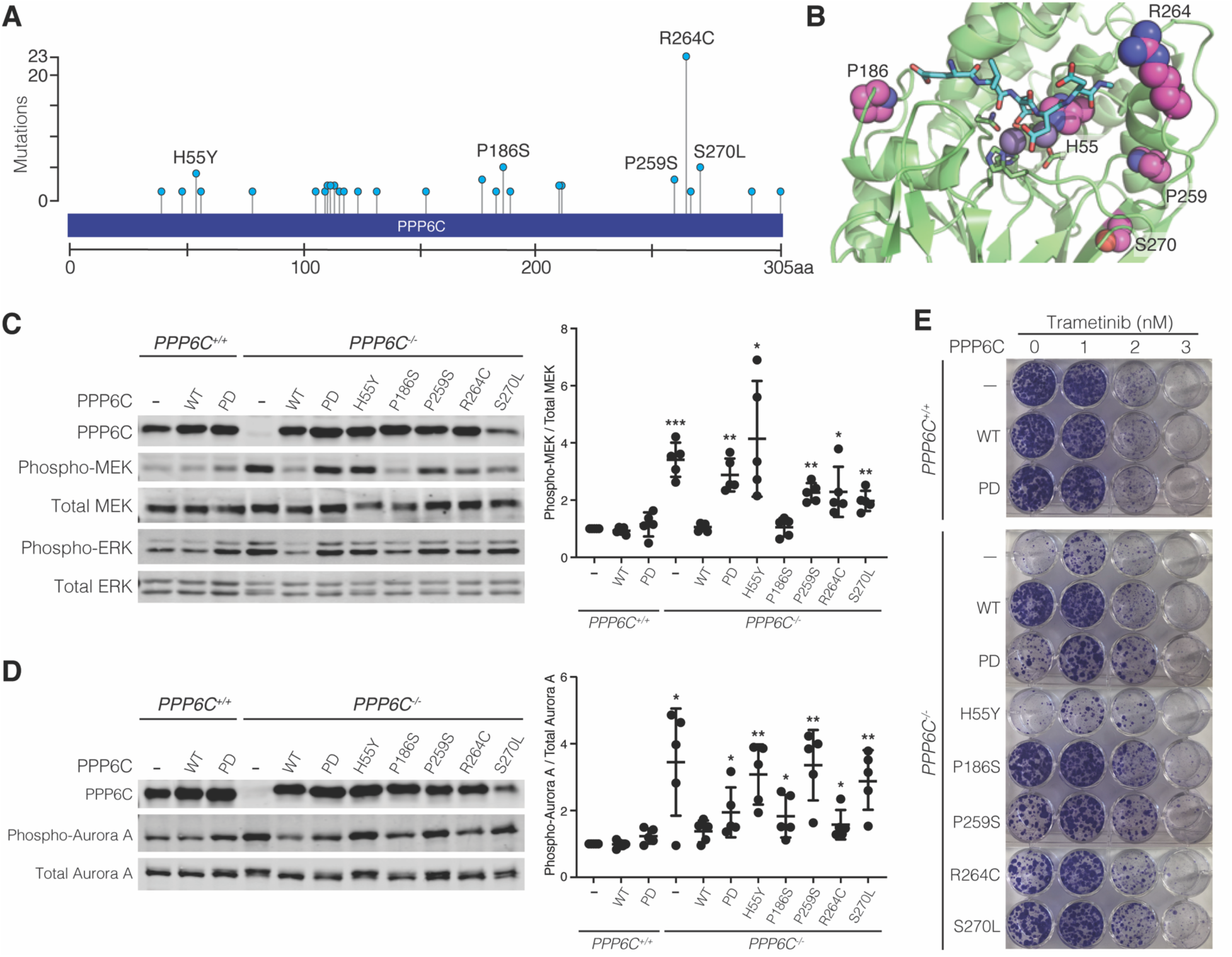
Cancer-associated PPP6C mutations abrogate PP6 phosphatase activity against MEK1/2. (A) Frequencies of PPP6C mutations reported in melanomas. Data are from nonoverlapping melanoma studies in cBioportal (Cerami et al., 2012; J. Gao et al., 2013). (B) PPP6C residues mutated in cancer are shown in spacefill representation modeled on the X-ray crystal structure of PPP5C in complex with a peptide substrate (PDB: 5HPE). The bound peptide is shown in cyan in stick representation, and the catalytic metal ions are shown as gray spheres. (C) *PPP6C*^*+/+*^ and *PPP6C*^*-/-*^ 501mel cells were transduced to stably express GFP (-), WT PPP6C, or the indicated PPP6C mutants. Cells were lysed and assessed by immunoblot for phosphorylated and total MEK and ERK. Phospho-MEK/total MEK signal ratios were quantified and normalized to the GFP-expressing *PPP6C*^*+/+*^ samples (*n* = 5). Significance is shown in comparison to *PPP6C*^*+/+*^ cells expressing GFP. *p < 0.05, **p < 0.01, ***p<0.001, paired t test. (D) Cells from (C) were treated with 100ng/mL nocodazole for 24 hours. Mitotic cells were lysed and assessed by immunoblot for phosphorylated and total Aurora A. Phospho-Aurora A/Total Aurora A signal ratios were quantified (*n* = 5) and significance determined as in (C). *p < 0.05, **p < 0.01, paired t test. (E) Cells from (C) were cultured in media containing DMSO vehicle alone or the indicated trametinib concentration for 2 weeks in colony forming assays. Colonies were stained with crystal violet.

We chose to characterize five of the most common mutations reported in melanomas (H55Y, P186S, P259S, R264C, and S270L) for their ability to regulate ERK signaling by ectopic expression in the *PPP6C*^-/-^ 501mel cell line (Figure 6A). While re-expression of WT PPP6C suppressed MEK and ERK phosphorylation to levels observed in parental cells, melanoma-associated PPP6C mutants varied in their impact on MEK phosphorylation (Figure 6C). In keeping with an essential role for His55 in catalysis, cells expressing the H55Y mutant exhibited the highest level of MEK phosphorylation, similar to that seen in empty vector control cells. Cells expressing the P259S, R264C, and S270L mutants had moderate but significant increases in MEK phosphorylation compared to cells expressing WT PPP6C, suggesting partial loss of activity. We note that the S270L mutant consistently expressed to lower levels than WT PPP6C or the other mutants, likely underlying its inability to promote MEK dephosphorylation. Ser270 maps to the globular core of the PPP6C catalytic domain (Figure 6C), potentially explaining the instability of the S270L mutation. Unlike the other mutants, expression of PPP6C^P186S^ reduced MEK phosphorylation to a similar extent as did the WT phosphatase. PPP6C mutants likewise impacted, to varying degrees, Aurora A phosphorylation in mitotically-arrested cells (Figure 6D). Expression of the H55Y, P259S, and S270L mutants resulted in high levels of Aurora A phosphorylation, similar to those seen in *PPP6C*^*-/-*^ cells. Aurora A phosphorylation was more modest in cells expressing the PD, P186S, and R264C mutants, but still significantly higher phosphorylation than in cells expressing WT PPP6C.

We next performed clonogenic assays to examine the impact of PPP6C mutation on cell growth and MEKi sensitivity (Figure 6E). We found that growth of the cells in the absence of drug inversely correlated with the degree of ERK pathway activation. For example, cells expressing the PPP6C^H55Y^ mutant, which had the highest levels of MEK and ERK phosphorylation, grew equivalently to PPP6C^-/-^ cells. Conversely, expression of PPP6C^P186S^, fully rescued the growth defect of null cells, in keeping with its complete reversal of MEK hyperactivation. The other mutants, which partially impacted MEK and ERK phosphorylation, likewise grew at an intermediate rate. In all cases, treatment with low concentrations of MEKi at least partially reversed the growth impairment observed in cells expressing PPP6C mutants, consistent with the inhibitor addiction phenotype observed with null cells. Collectively, these experiments indicate that cancer-associated PPP6C mutations generally cause partial loss-of-function, impacting both dephosphorylation of substrates and sensitivity to MEKi.

## DISCUSSION

PPP6C is thought to be a tumor suppressor in melanoma (Cancer Genome Atlas Network, 2015; Hodis et al., 2012; Krauthammer et al., 2015; Malicherova et al., 2019; Palmieri et al., 2018), yet our understanding of how it modulates cancer relevant pathways is limited. PPP6C has well-established roles in mitotic progression through dephosphorylation of the kinase Aurora A (Zeng et al., 2010), DNA-dependent protein kinase (DNA-PK) (Douglas et al., 2010; Hosing et al., 2012), the condensin subunit NCAP-G (Rusin et al., 2015), as well as other substrates. Most studies of PPP6C in melanoma have focused on its regulation of Aurora A, an essential kinase regulating mitotic spindle assembly and chromosome segregation (Hammond et al., 2013; Zeng et al., 2010). Melanoma-associated PPP6C mutations impair its ability to dephosphorylate and inactivate Aurora A, resulting in genomic instability and DNA damage that may be an early event contributing to cancer progression (Gold et al., 2014; Hammond et al., 2013). Our identification of PPP6C as a MEK phosphatase suggests that it also acts as a negative regulator of the core pathway driving melanoma, likely underlying at least in part its role as a tumor suppressor. Impaired dephosphorylation of MEK may also be relevant to previous studies showing that loss of PPP6C promotes oncogenic RAS driven tumors in mouse keratinocytes (Kurosawa et al., 2018) and in *Drosophila* (Ma et al., 2017). PPP6C mutations in melanoma almost exclusively co-occur with BRAF and NRAS mutations, suggesting that alone they do not provide oncogenic levels of ERK signaling. In this context, downregulation of PPP6C is likely to have a role in tuning flux through the ERK pathway to counteract negative feedback regulation. A similar phenomenon may drive selection for mutations in MEK1, MEK2 and ERK2 found at low frequency in melanomas that generally co-occur with other activating lesions (Cancer Genome Atlas Network, 2015; Y. Gao et al., 2018; Hodis et al., 2012; Palmieri et al., 2018).

The observation that loss or mutation of PPP6C is deleterious to cell growth may appear at odds with its role as a negative regulator of ERK signaling. Recent studies indicate however that in the context of activating BRAF and RAS mutations, further elevation of signaling through the ERK pathway is toxic (Hong et al., 2018; Kong et al., 2017; Leung et al., 2019; Moriceau et al., 2015). This phenomenon gives rise to inhibitor addiction, in which tumor cells treated with pathway inhibitors reactivate ERK signaling to re-establish signaling within an optimal range, or “fitness zone”. Subsequent inhibitor withdrawal results in a rebound signaling, leading to toxic hyperactive signaling outside of the fitness zone. BRAFi and/or MEKi addicted melanoma tumors grown in mice regress when treatment with inhibitors is ceased (Hong et al., 2018; Kong et al., 2017), suggesting periodic “drug holidays” could benefit patients who have progressed on BRAFi and MEKi. However, in clinical studies, cessation of BRAFi and/or MEKi therapy for several months re-sensitized to the inhibitors, though did not cause tumor regression (Rogiers et al., 2017; Seghers et al., 2012), revealing how the diversity of resistance mechanisms, tumor heterogeneity and adaptability complicate response to drug withdrawal in patients. Our studies suggest downregulation or inactivation of PPP6C as an unappreciated mechanism contributing to inhibitor addiction and resistance. This substantiates the identification of PPP6C as a candidate BRAFi resistance gene in an insertional mutagenesis screen in a mouse model of melanoma, and as a MEKi resistance gene in CRISPR/Cas9 screens conducted in NRAS mutant melanoma cells in culture (Hayes et al., 2019; Perna et al., 2015).

The toxicity associated with high level ERK signaling indicates that tumor cells harboring hyperactivating BRAF or RAS mutations rely on negative feedback control to maintain signaling within the fitness zone. For example, silencing expression of the ERK phosphatase DUSP6 is toxic to BRAF mutant melanoma and KRAS mutant lung cancer cells (Unni et al., 2018; Wittig-Blaich et al., 2017). Indeed, in genome wide CRISPR/Cas9 screens, dependency on DUSP6 was highly correlated with dependency on PPP6C across a panel of melanoma cells lines, in keeping with PPP6C as a key negative regulator of ERK signaling. We hypothesize that through dephosphorylation of MEK, PPP6C likewise contributes to negative feedback control of the ERK pathway. While we did not observe changes in levels of PPP6C upon inhibition of BRAF-MEK-ERK signaling, we cannot rule out transcriptional control of PP6 regulatory or scaffolding subunits as a mechanism of feedback regulation. Furthermore, because phosphorylation of PP6 regulatory subunits can mediate recruitment to substrates and other interaction partners, targeting of MEK may be impacted by ERK-dependent phosphorylation or other modifications.

Atypically for a tumor suppressor, more than half of melanoma-associated PPP6C mutations occur recurrently in hotspots (Figure 6A), while frameshift/truncation mutations are relatively uncommon. Notably, non-recurrent mutations, while distributed in the primary sequence, do significantly cluster in the phosphatase catalytic center. Prior analysis of recurrent and non-recurrent PPP6C mutations indicate that all of them, to varying degrees, reduce catalytic activity (Gold et al., 2014; Hammond et al., 2013). These prior results are consistent with our observation that PPP6C mutations vary in their ability to suppress MEK phosphorylation. Given that PPP6C has been characterized as a “common essential” gene, it is possible that full loss of function is incompatible with cell proliferation. It has also been reported that PPP6C mutations weaken association with the PPP6R2 regulatory and ANKRD28 scaffolding subunits (Hammond et al., 2013). While the three-dimensional structure of a PP6 heterotrimer has not yet been determined, in the X-ray crystal structures of the PP2A-B56 holoenzyme, the residue analogous to R264 participates in interactions between catalytic and regulatory subunit (Y. Xu et al., 2006). In contrast, the same residue is not at the catalytic-regulatory subunit interface in structures of other PP2A holoenzymes (Wlodarchak et al., 2013; Y. Xu et al., 2008). This raises the possibility that PPP6C mutations might change the heterotrimer composition, favoring some regulatory subunits over others, which could favor selective dephosphorylation of substrates in a manner that preserves cell viability. By analogy, recurrent cancer-associated mutations in the PP2A scaffolding subunit PPP2R1A preferentially disrupt interactions with some regulatory B subunits rather than causing complete loss of function (O’Connor et al., 2020). The capacity for specific complexes to restrain cell proliferation is key to the activity of recently developed PP2A small molecule activators, which stabilize specific holoenzymes (Leonard et al., 2020; Morita et al., 2020). As with PP2A, at least some PP6 substrates are recruited through interaction with individual regulatory subunits (Heo et al., 2020; Hosing et al., 2012; Stefansson and Brautigan, 2006; Zhong et al., 2011), suggesting the potential for developing PP6 activators with therapeutic benefit.

While our studies implicate PP6 as a MEK phosphatase, early reports suggested that MEK is dephosphorylated by PP2A (Gómez and Cohen, 1991; Sontag et al., 1993). Indeed, downregulation of PP2A activity by loss or mutation of its scaffolding subunit PPP2R1A causes resistance to MEK inhibitors in KRAS mutated lung and colorectal cancer cell lines (Kauko et al., 2018; O’Connor et al., 2020). Likewise, enhancing PP2A activity through downregulation of endogenous inhibitor proteins or through small molecule activators sensitizes to MEK inhibition (Kauko et al., 2018). However, in these contexts PP2A promotes sensitivity to MEKi by restraining bypass PI3K/mTOR signaling and by direct dephosphorylation of MYC or ERK itself. Likewise, the tumor suppressor function of PP2A is suggested to involve other processes, such controlling the stability of MYC and β-catenin (Sablina et al., 2010; Yeh et al., 2004). Notably, in addition to restraining ERK signaling, PP2A can also promote MEK phosphorylation through dephosphorylation of inhibitory feedback phosphorylation sites on RAF and KSR (Ory et al., 2003). Because oncogenic mutant BRAF signals independently of KSR and is feedback-resistant (Ritt et al., 2010), PP2A activity as a MEK phosphatase would not be counterbalanced by activation of upstream signaling. This phenomenon may explain our observation that like PP6, PP2A also contributes to MEK dephosphorylation in BRAF mutant 501mel cells. While we found that silencing PPP6C expression hyperactivates MEK in most cell lines we examined, this was not universally the case. The relative contributions of PP2A, PP6 and potentially other phosphatases to dephosphorylation of MEK is thus context dependent. Melanoma cells in particular are characterized by low PP2A activity, and the PP2A inhibitor protein CIP2A is an established transcriptional target of ERK signaling (Khanna et al., 2011; Mannava et al., 2012). Interestingly, we observed that PP6 and PP2A catalytic subunit expression levels increase when the other is silenced (Figure 5E), suggesting compensation between the two complexes that could potentially influence dephosphorylation of any number of targets. Further work will be necessary to understand the lineage-specific, signaling, or genetic contexts that dictate PPP6C regulation of MEK1/2.

## ACKNOWLEDGMENTS

We thank David Stern for helpful comments on the manuscript, and Valia Mihaylova for assistance with the shRNA screen. We are grateful to Ruth Halaban and Antonietta Bacchiocchi for providing cell lines from the Yale SPORE in Skin Cancer Biospecimen Core, and to Harriet Kluger, Craig Crews, and Narendra Wajapeyee for providing additional cell lines. We thank Titus Boggon, Derek Abbott, and Natalie Ahn for providing plasmids. This work was supported by NIH R01 GM102262 and a developmental research project from the Yale SPORE in Skin Cancer (NIH P50 CA121974) to B.E.T. Support for E.C. was provided by a Ruth L. Kirschstein Predoctoral National Research Service Award (NIH F31 CA220999) and Institutional National Research Service Award (T32 GM007324).

## AUTHOR CONTRIBUTIONS

Conceptualization, E.C. and B.E.T.; Methodology, E.C. and B.E.T.; Investigation, E.C. and H.J.L.; Formal Analysis, E.C. and L.K.; Writing-Original Draft, E.C. and B.E.T.; Writing-Review & editing, E.C., B.E.T., D.A.C.; Visualization, E.C. and B.E.T; Funding Acquisition E.C. and B.E.T.; Resources, E.C. B.E.T, and D.A.C.; Supervision, E.C., B.E.T and D.A.C.

## DECLARATION OF INTERESTS

The authors declare no competing interests.

## STAR Methods

### KEY RESOURCES TABLE

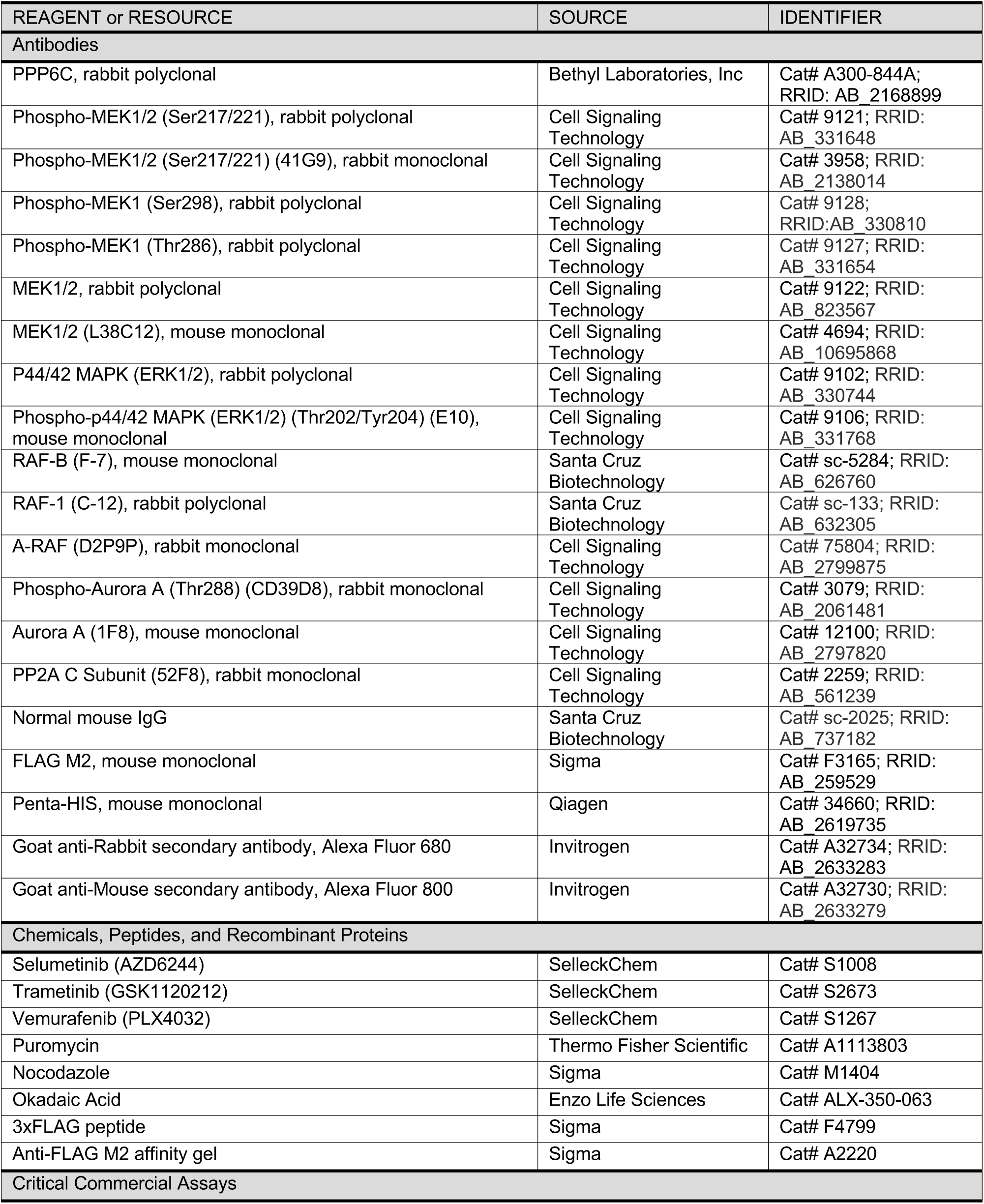
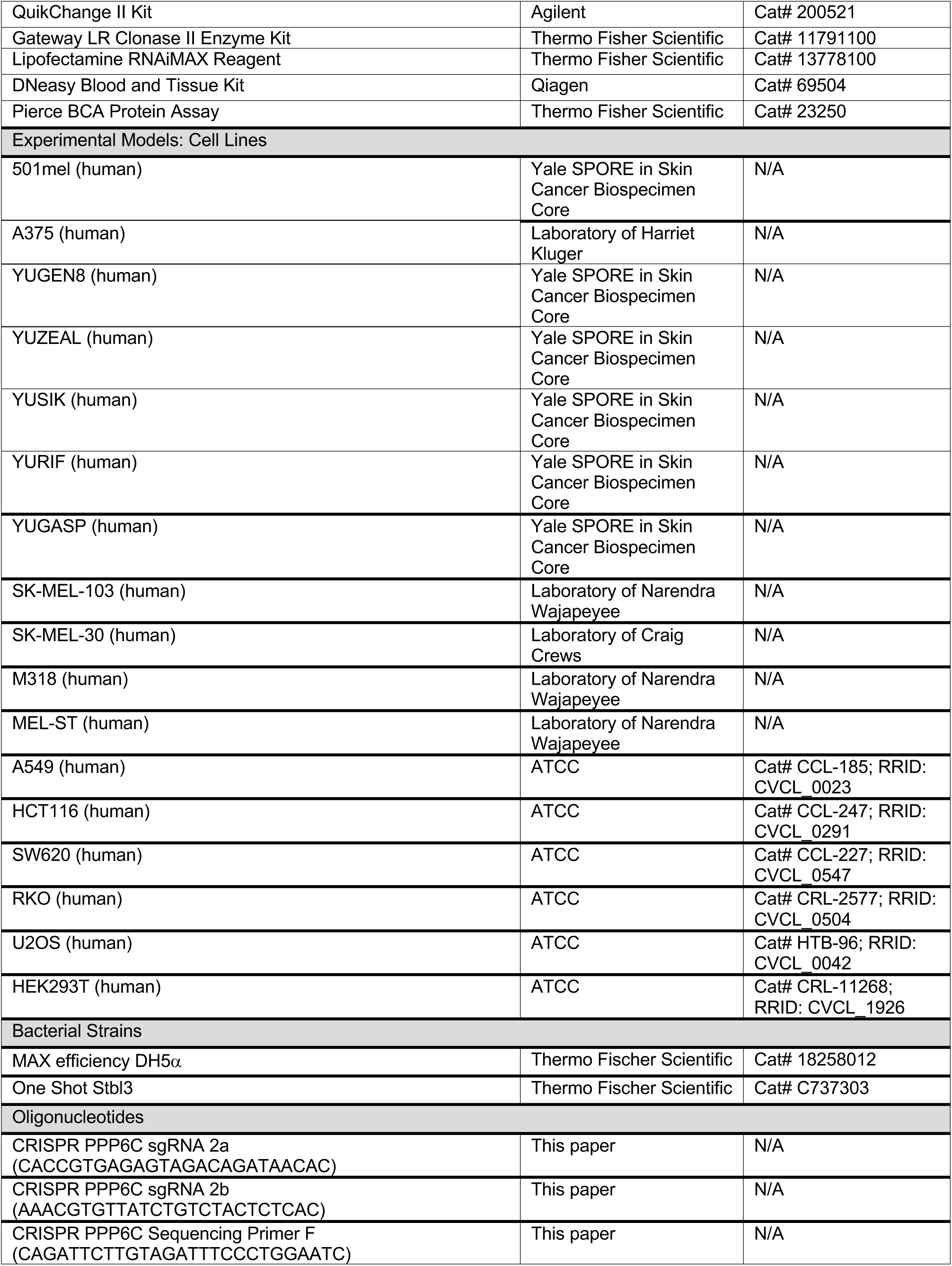
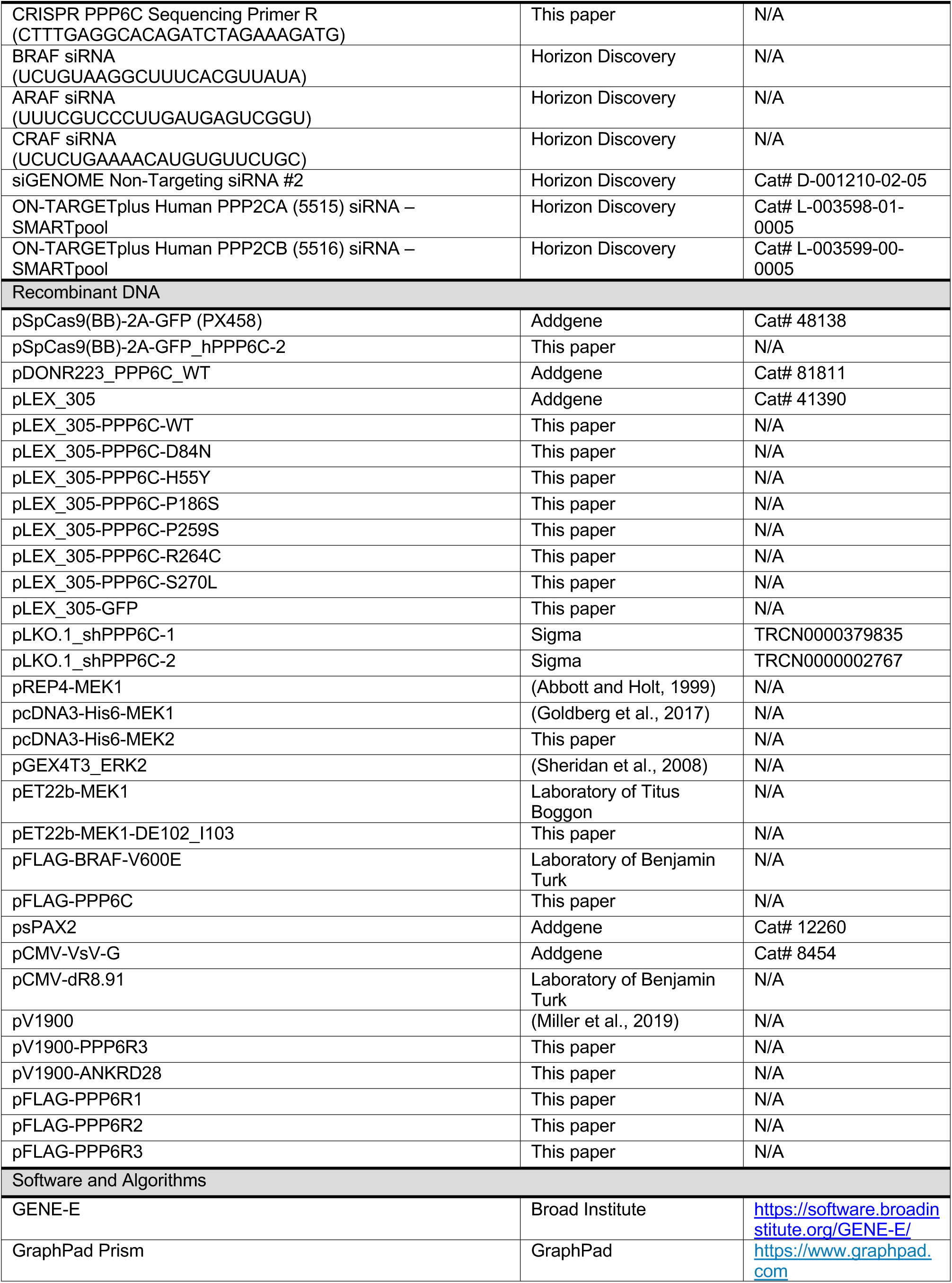
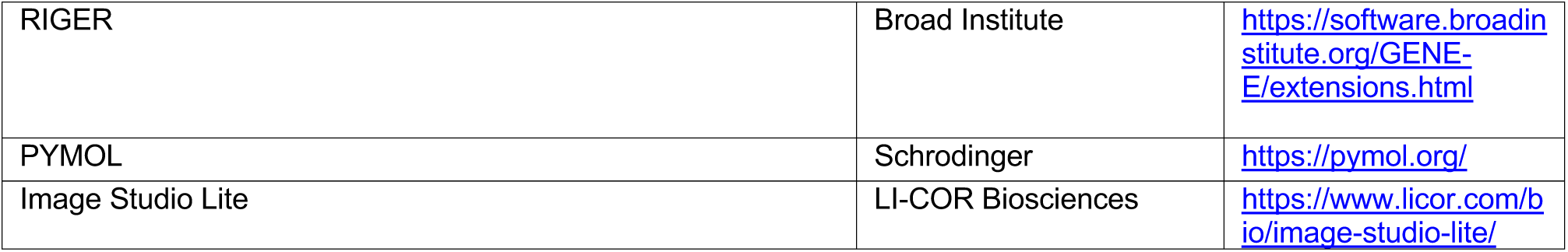

### RESOURCE AVAILABILITY

Further information and requests for resources and reagents should be directed to and will be fulfilled by the lead contact, Ben Turk.

### EXPERIMENTAL MODEL AND SUBJECT DETAILS

#### Cell Lines and Culture Conditions

501mel, YUGEN8, YUZEAL, YUSIK, YURIF, and YUGASP cells were cultured in Opti-MEM medium (Gibco) supplemented with 5% fetal bovine serum (FBS) (Gibco) and 1% penicillin/streptomycin (P/S, Gibco). A375 cells were cultured in Opti-MEM medium supplemented with 10% FBS and 1% P/S. MEL-ST, U2OS, and HEK293T cells were cultured in DMEM medium (Gibco) supplemented with 10% FBS and 1% P/S. SK-MEL-103, SK-MEL-30, HCT116, and M318 cells were cultured in RPMI 1640 medium (Gibco) supplemented with 10% FBS and 1% P/S. RKO were cultured in MEM medium (Gibco) supplemented with 10% FBS and 1% P/S. All cell lines were cultured at 37°C under 5% CO_2_.

### METHOD DETAILS

#### Plasmids, Cloning, and Mutagenesis

Plasmids harboring cDNAs of PPP6C, PPP6R1, PPP6R3 and ANKRD28 in pDONR223 were from the human ORFeome collection (v8.1), and the PPP6R2 cDNA was from Transomic. The lentiviral expression vector pLEX_305-PPP6C and the transient expression plasmid pVL1900-ANKRD28 (untagged) were generated by Gateway recombination cloning into their respective destination vectors. The untagged expression vector for PPP6R3 used for preparation of PP6 complexes was generated by Gateway recombination into pV1900 followed by QuikChange mutagenesis to insert a stop codon upstream of the FLAG tag. The transient expression vector for N-terminally FLAG-tagged PPP6C, PPP6R1, PPP6R2 and PPP6R3 were made by PCR amplification of the coding sequence from the source plasmid and inserting into pcDNA3-FLAG by either Gibson assembly (PPP6R3) or restriction enzyme cloning (all others). The mammalian expression vector for N-terminally 6xHis-tagged MEK2 was generated by shuttling the entire coding sequence from pRSET-MEK2 (obtained from the laboratory of Natalie Ahn) and into pcDNA3. All mutants were generated using QuikChange Site Directed Mutagenesis following standard protocols. Constructs were verified by Sanger sequencing through the entire open reading frame.

#### Recombinant Lentivirus Production and Cell Infection

shRNA lentiviruses were packaged in low passage HEK293T cells by polyethylenimine (PEI) co-transfection with packaging constructs dR8.91 and VsV-G (Addgene, 8454). PPP6C expression lentiviruses were packaged in low passage 293T cells by PEI co-transfection with packaging constructs psPAX2. (Addgene, 12260) and VsV-G (Addgene, 8454). For PEI co-transfection, lentiviral transfer plasmid:packaging plasmid:envelope plasmid ratio was at 10:10:1 with the PEI:DNA ratio at 3:1. Supernatant media containing virus was collected at 48 hours post transfection. Cells were infected with lentivirus at an MOI of 0.3-0.4 in the presence of 4ug/mL polybrene for 24 hours and selected for >48 hours in fresh media containing (1.5-2.5 ug/mL) puromycin.

#### shRNA Screening

The shRNA library was custom generated by pooling human MISSION shRNA constructs (Sigma) targeting all annotated protein kinases and phosphatases (Table S2) and packaged into lentiviral particles as described above. To initiate the screen, 501mel cells were transduced for 24 hours with the lentiviral library in 0.4 μg/ml polybrene at an MOI of 0.3 to assure that most cells receive a single viral integration. A sufficient number (8 × 10^6^) of cells were infected to ensure >1,000-fold coverage for each unique shRNA in the library for a reference sequencing sample and for each drug condition. Infected cells were selected with 1.8 μg/mL puromycin for 48 hours, trypsinized, and 8 × 10^6^ cells were reserved for the T_0_ reference sample. For the remainder, 8 × 10^6^ cells were plated for each of the 5 conditions: 0.0001% DMSO vehicle control, 1 nM trametinib, 3.3 nM trametinib, 33 nM selumetinib, and 100 nM selumetinib. Every two doublings, cells were counted, and 8 × 10^6^ cells were replated for propagation. The screen was carried out for 10 total population doublings (T_10_). Genomic DNA from the T_0_ and T_10_ samples for each of the drug conditions was extracted using Qiagen DNeasy Blood and Tissue Kit (Qiagen, Cat No. 69504), following the manufacturer’s protocol. For each drug condition/time point sample, the shRNA integrants were PCR-amplified from the genomic DNA with barcoded primers and sequenced on an Illumina HiSeq instrument. The RIGER algorithm in GENE-E (www.broadinstitute.org/cancer/software/GENE-E/) was used to rank each gene by their enrichment.

#### MEKi Dose Response Assays

Cells (750 per well) were seeded in 96-well black/clear bottom plates, allowed to recover overnight, and treated with varying concentrations of trametinib or selumetinib (6 wells/concentration) in fresh media for 72 h. Media aspirated and replaced with fresh media containing 44 μM resazurin (alamarBlue Cell Viability Reagent, Fisher Scientific). Plates were incubated in the dark for 4 hours at 37°C, and fluorescence (excitation 560 nm; emission 590nm) was measured on a plate reader. When MEKi treatment was initiated, starting time reading was obtained on a separate plate containing untreated cells. Starting point readings were subtracted from the 72 h readings to measure overall growth inhibition. Dose response curves were generated with GraphPad Prism.

#### Clonogenic Growth Assays

Cells (1 × 10^3^) were plated in each well of a six-well plate containing 3 mL of media with or without MEKi and cultured at 37°C under 5% CO_2_ undisturbed for 14 days. Media was removed, and cells were gently washed with PBS. The cells were stained with crystal violet staining solution (0.5% crystal violet, 6% formaldehyde, 1% methanol in PBS) for 15 min and washed 3 times with water. Plates were air-dried and imaged. For experiments characterizing PPP6C mutants, cells (2.5 × 10^3^) were plated in 12 well plates containing 1 mL media with or without MEKi.

#### Cell Lysis and Immunoblot Analysis

Cells were placed on ice, washed twice with cold PBS, and lysed in cold lysis buffer (20 mM Tris [pH 8.0], 137 mM NaCl, 10% glycerol, 1% Igepal CA-630, 1 mM PMSF, 1 mM Na_3_VO_4_, 10 μg/mL leupeptin, 2 μg/mL pepstatin A, 10 μg/mL aprotinin) for 15 min. Cell lysates were scraped into 1.5 mL tubes and clarified in a 4 °C microcentrifuge at 13,000 rpm for 10 min. Cleared lysates were analyzed by BCA protein assay. 4X SDS-PAGE loading buffer was added to lysates to prepare immunoblot samples. Equal amounts of lysate (15 μg per lane) were fractionated by SDS-PAGE and transferred to polyvinyl difluoride (PVDF) (Sigma, IPFL85R) membrane. Membranes were blocked in Tris buffered saline (TBS) with 5% non-fat milk for one h and probed overnight at 4 °C with primary antibodies diluted according to manufacturer’s recommendations. Membranes were incubated for 1 h in fluorescently-labeled secondary antibodies diluted 1:10,000 in TBS with 5% bovine serum albumin (BSA) and 0.1% Tween20. Western blots were imaged with an Odyssey CLx imaging system (LI-COR Biosciences), and densitometry-based quantification was carried out with Image Studio Lite.

#### Co-immunoprecipitation

HEK293T cells in 10 cm plates were transiently transfected with equal amounts of pREP-MEK1 (untagged) and FLAG tagged PP6 subunit plasmid precomplexed with polyethyleneimine (PEI) at a 3:1 ratio with DNA. After 48 h, cells were placed on ice and washed twice with cold PBS. On the second PBS wash, cells were scraped into 1.5 mL tubes and pelleted at 1000 rpm for 5 min. Cells were resuspended in 300 μL hypotonic lysis buffer (10 mM Tris-HCl [pH 8.0], 1 mM KCl, 1.5 mM MgCl_2_, 0.5 mM DTT, 0.05% Igepal CA-630, 1 mM PMSF, 1 mM Na_3_VO_4_, 10 μg/mL leupeptin, 2 μg/mL pepstatin A, 10 μg/mL aprotinin) and kept on ice for 5 min. Cell lysates were vortexed for 1 minute and run three times through a 25G needle with a syringe. Lysates were spun at 3500 rpm for 10 minutes in a 4°C microcentrifuge. A portion (30 μL) of the supernatant was reserved for analysis of the whole cell lysate sample. The remaining total supernatant was brought to a volume of 500 μL with additional hypotonic lysis buffer, and the [NaCl] was adjusted to 150 mM. Anti-FLAG M2 Affinity Gel beads (Sigma, A2220) were blocked in 5% BSA-TBS solution for 1 h, equilibrated to hypotonic lysis buffer, and 30 μL of the suspension was added to each supernatant. Samples were rotated at 4°C overnight. Beads were pelleted and washed with cold wash buffer 1 (20 mM Tris [pH 7.5], 150 mM NaCl, 1% Triton X-100, 2.5 mM Na_4_P_2_O_7_, 1 mM β-glycerophosphate, 3 mM β-mercaptoethanol, 1 mM PMSF, 1 mM Na_3_VO_4_, 10 μg/mL leupeptin, 2 μg/mL pepstatin A, 10 μg/mL aprotinin) for 10 minutes, followed by one quick and one 10 min wash with cold wash buffer 2 (50 mM HEPES [pH 7.4], 150 mM NaCl, 3 mM 2-mercaptoethanol, 0.1 mM Na_3_VO_4_, 0.01% Igepal CA-630, 10% glycerol). Beads were resuspended in 30 μL 2X SDS buffer (100mM Tris-Cl [pH 6.8], 4% SDS, 20% glycerol) and boiled for 5 min. Samples were centrifuged in Whatman UNIFILTER 0.45 μm plates (Sigma, WHA77002808) at 4000 rpm for 10 minutes to remove beads, and 4X SDS-PAGE loading buffer was added to filtrates. Samples were analyzed via immunoblot as described above.

#### Generation of CRISPR/Cas9 Knockout Cell Lines

CRISPR/Cas9 constructs were generated by cloning sgRNA sequences into pSpCas9(BB)-2A-GFP (Addgene, 48138) according to the cloning protocol established by the Zhang lab (https://www.addgene.org/browse/article/7475/). Two sets of sgRNA oligos were used but only one sgRNA targeting exon 4 resulted in PPP6C knockout clones. pSpCas9(BB)-2A-GFP was used to generate negative control clones. 501mel cells were transfected with CRISPR/Cas9 constructs via PEI, and 48 hours post transfection, cells were trypsinized. After centrifugation and removal of media/trypsin, cells were resuspended in PBS and transferred to FACS tubes. Single GFP positive cells were sorted into 96 well plates via a BD FACSAria instrument. 96 well plates were treated with 0, 1, or 2 nM trametinib and incubated for several weeks until colonies were observed. PPP6C knockout 501mel cell colonies only grew out in the presence of trametinib and were maintained in 1-2 nM trametinib but withdrawn from trametinib >24 h before experiments. PPP6C knockout was confirmed by immunoblot and sanger sequencing of PCR amplified target site.

#### Protein Purification

MEK1 was expressed in HEK293T cells by PEI transfection with pcDNA3-His-MEK1 alone or in a 4:1 ratio with pFLAG-BRAF-V600E to generate phosphorylated MEK1. After 40 h, plates were put on ice and washed once with ice-cold PBS. To lyse cells, 1 mL ice cold lysis buffer (20 mM Tris [pH 7.5], 150 mM NaCl, 1% Triton X-100, 2.5 mM Na_4_P_2_O_7_, 1 mM β-glycerophosphate, 1mM Na_3_VO_4_, 3 mM β-mercaptoethanol, 1 mM PMSF, 10 μg/mL leupeptin, 2 μg/mL pepstatin A, 10 μg/mL aprotinin) was added to each plate. Lysates were scraped into 1.5 mL tubes, incubated on ice for 10 minutes, and clarified in a microcentrifuge at 13,000 rpm for 10 min at 4°C. Supernatants were transferred to fresh tubes, and 50 μL of Talon resin (Takara) was added. Samples were rotated for 2 hours at 4 °C. Beads were pelleted for (2 min, 4000 rpm) at 4°C microcentrifuge, washed twice with lysis buffer containing 10 mM imidazole, and transferred into a column. Beads were washed with 2 mL of wash buffer (50 mM HEPES [pH 7.4], 150 mM NaCl, 3 mM 2-mercaptoethanol, 10 mM imidazole, 0.01% Igepal CA-630, 10% glycerol), and MEK1 was eluted in 150 μL fractions with wash buffer + 250 mM imidazole. The two most concentrated fractions as determined by Bradford assay (Bio-Rad, 5000006) were combined and dialyzed overnight at 4°C into 20 mM HEPES [pH 7.4], 150 mM NaCl, 1 mM DTT, 10% glycerol, 0.01% Igepal CA-630. Protein concentration was estimated from Coomassie-stained 10% polyacrylamide gels using a BSA standard curve

To prepare PP6 complexes, HEK293T cells were co-transfected in 15 cm plates with 4 μg pFLAG-PPP6C, 8.6 μg pV1900-PPP6R3 and 8.6 μg pV1900-ANKRD28 pre-complexed with 63.3 μg PEI. Cells were lysed 40 hours post-transfection after washing with cold PBS in 2.25 μL CHAPS lysis buffer (50 mM Tris [pH 7.5], 150 mM NaCl, 0.3% CHAPS, 1 mM PMSF, 10 μg/mL leupeptin, 2 μg/mL pepstatin A, 10 μg/mL aprotinin) per plate. Lysates were cleared as above, and M2 anti-FLAG affinity gel (33 μL per plate) was added to the supernatant. Samples were rotated at 4 °C for 1 hr, and beads were pelleted, washed three times with lysis buffer (0.3 mL per plate) and once with wash buffer (50 mM HEPES, pH 7.4, 150 mM NaCl, 10% glycerol). Protein was eluted in two rounds with 30 uL wash buffer + 0.5 mg/mL 3xFLAG peptide (Sigma F4799) per plate, snap frozen on dry ice/EtOH and stored at −80 °C. Protein concentration was estimated from Coomassie-stained 10% polyacrylamide gels using a BSA standard curve.

ERK2 was purified in unphosphorylated from bacteria as described in (Sheridan et al., 2008). ERK2 (21 μM) was phosphorylated *in vitro* by incubation with 0.2 μM bacterially-expressed active His_6_-MEK1^ΔE102-I103^ in kinase reaction buffer (50 mM Tris [pH 8.0], 50 mM NaCl, 0.5 mM ATP, 1mM DTT, 0.01% Igepal CA-630, 10% glycerol, 10 mM MgCl_2_) at 30°C for 30 min. MEK1 was removed by adding 20 μL Talon resin, rotating at 4°C for 1 h, and filtered through a chromatography column.

#### BRAF IP Kinase Assay

Protocol for BRAF IP kinase assays was adapted from (Bondzi et al., 2000). Confluent 10 cm plates of shCTRL, shPPP6C-1, or shPPP6C-2 expressing 501mel cells were washed twice with cold PBS and lysed in RIPA buffer (20 mM Tris [pH 8.0], 137 mM NaCl, 10% glycerol, 1% NP-40, 1 mM PMSF, 1 mM Na_3_VO_4_, 10 μg/mL leupeptin, 2 μg/mL pepstatin A, 10 μg/mL aprotinin, 0.1% SDS, 0.5% sodium deoxycholate) on ice for 15 min. Cells were scraped into 1.5 mL tubes and lysates were passed through 22G needle with a syringe 3 times. Lysates were clarified in a 4°C microcentrifuge at 13,000 rpm for 10 min. Lysates were analyzed by BCA protein assay and equivalent amounts of protein were pre-cleared for 1 hour at 4°C with nProtein A Sepharose 4 Fast Flow beads (Sigma, GE17-5280-01) pre-equilibrated with lysis buffer. Beads were removed and lysates were divided into 500 μL aliquots containing 500 μg protein for each assay condition or timepoint. Antibody to BRAF (7.5 μL) was added to each sample, and tubes were rotated 2 h at 4°C. To precipitate immune complexes, 50 μL of a 1:1 suspension of nProtein A Sepharose in lysis buffer was added, and tubes again rotated for 2 h at 4°C. Beads were pelleted, washed three times with lysis buffer, and resuspended in kinase reaction buffer (20 mM Tris [pH 7.4], 20 mM NaCl, 1 mM DTT, 10 mM MgCl_2_, 1 mM MnCl_2_). Purified unphosphorylated MEK1 (0.5 μg) and/or vemurafenib (to 1 μM) were added to as indicated. To initiate kinase reactions, ATP was added to a final concentration of 1 mM and volume of 40 μL, and tubes were transferred to 30°C heat block for the indicated times. Reactions were quenched with 4X SDS-PAGE loading buffer and boiled for 5 minutes and then subjected to SDS-PAGE (10% acrylamide) and immunoblotting as described above.

#### siRNA Transfection

Cells plated in 6-well plates were transfected with siRNA using Lipofectamine RNAiMAX reagent (Thermo Fisher Scientific, 13778100). Equal parts siRNA oligonucleotides (100 nM in 1X siRNA buffer, Horizon Discovery) and Lipofectamine RNAiMAX (diluted 1:100 in Opti-MEM Medium) were combined and incubated for 15 minutes at room temperature. Lipofectamine:siRNA complexes (400 μL) and complete media (600 μL) were added to each well. Cells were incubated for 72 h before being lysed for immunoblot analysis as described above. For cells transfected with two different siRNAs, half the amount of each siRNA was used.

#### In Vitro Phosphatase Assays

For each reaction, PP6 complex containing 125 ng PPP6C and 500 ng Phospho-MEK were mixed in 30 μL reaction buffer (50 mM Tris-HCl [pH8.0], 0.5 mM MnCl_2_, 2 mM DTT) with or without 100nM okadaic acid as indicated. Reactions were incubated at 30 °C for the indicated time, quenched by the addition of 4X SDS-PAGE loading buffer, and boiled for 5 min. Samples were separated by SDS-PAGE (10% acrylamide) and analyzed by immunoblot.

## Notes

### Competing Interest Statement

The authors have declared no competing interest.

