## Supplemental Information for "PPP6C negatively regulates oncogenic ERK signaling through dephosphorylation of MEK"

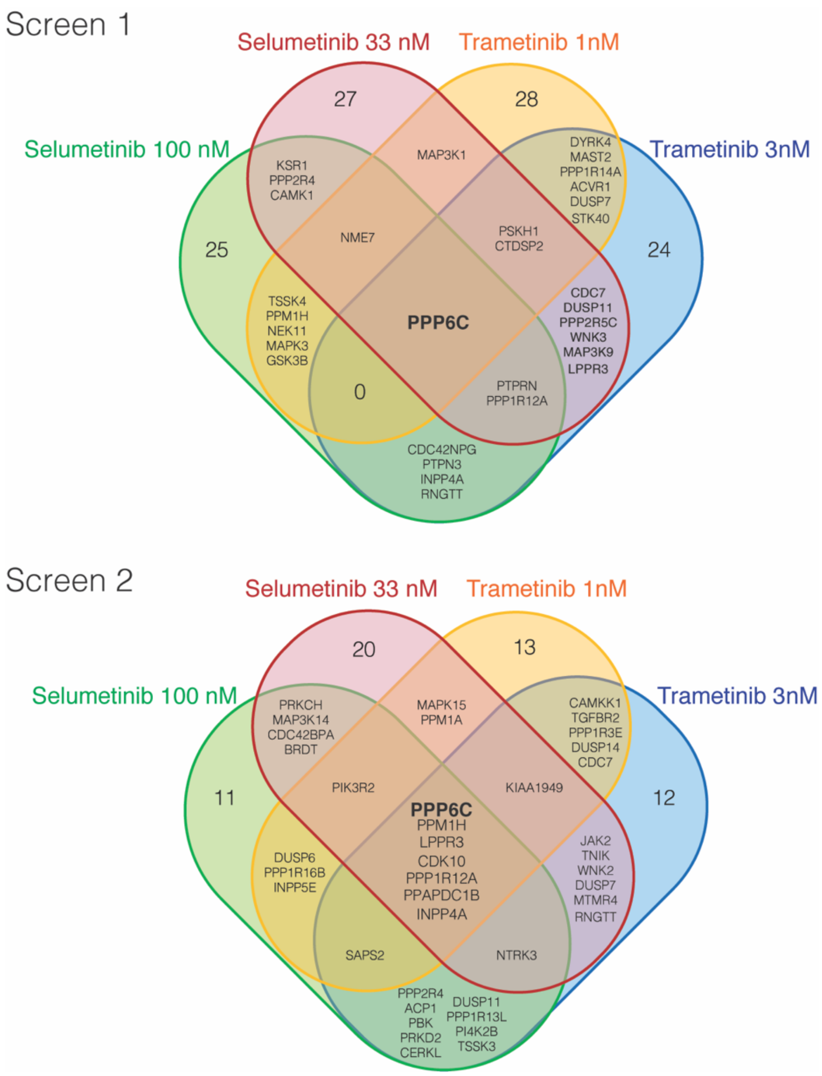


**Figure S1. Hits from MEKi sensitivity shRNA screens**
Venn diagrams of top 50 enriched genes for each drug condition.

**A**

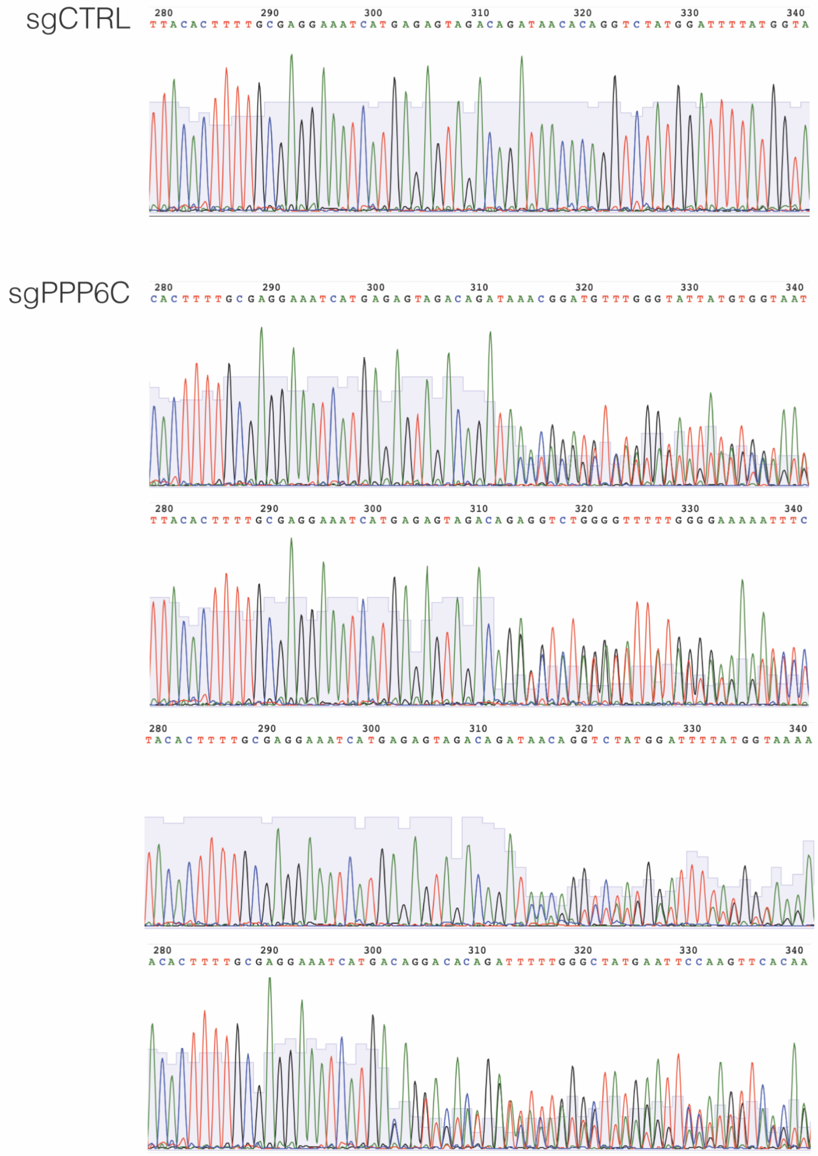
sgCTRL
sgPPP6C

**B**

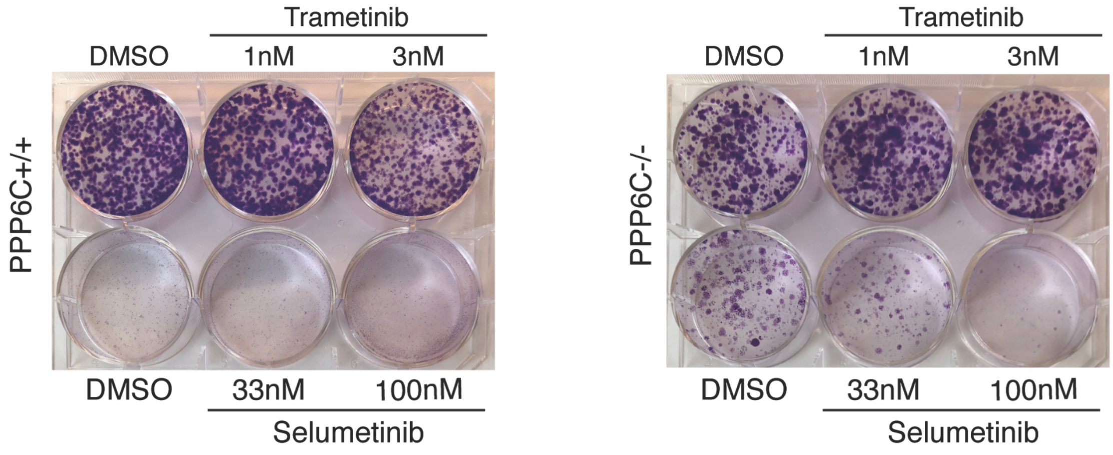
Trametinib
DMSO
1nM
3nM
PPP6C+/+
DMSO
33nM
100nM
Selumetinib
Trametinib
DMSO
1nM
3nM
PPP6C-/-
DMSO
33nM
100nM
Selumetinib

**Figure S2. PPP6C CRISPR/Cas9 knockout cell lines**
(A) Sanger sequencing chromatograms for *PPP6C*$^{+/+}$ and *PPP6C*$^{-/-}$ clonal 501mel cell lines generated by CRISPR/Cas9.
(B) *PPP6C*$^{+/+}$ and *PPP6C*$^{-/-}$ 501mel cells were cultured in media containing DMSO or the indicated concentration of trametinib for 2 weeks in colony forming assays. Colonies were stained with crystal violet.

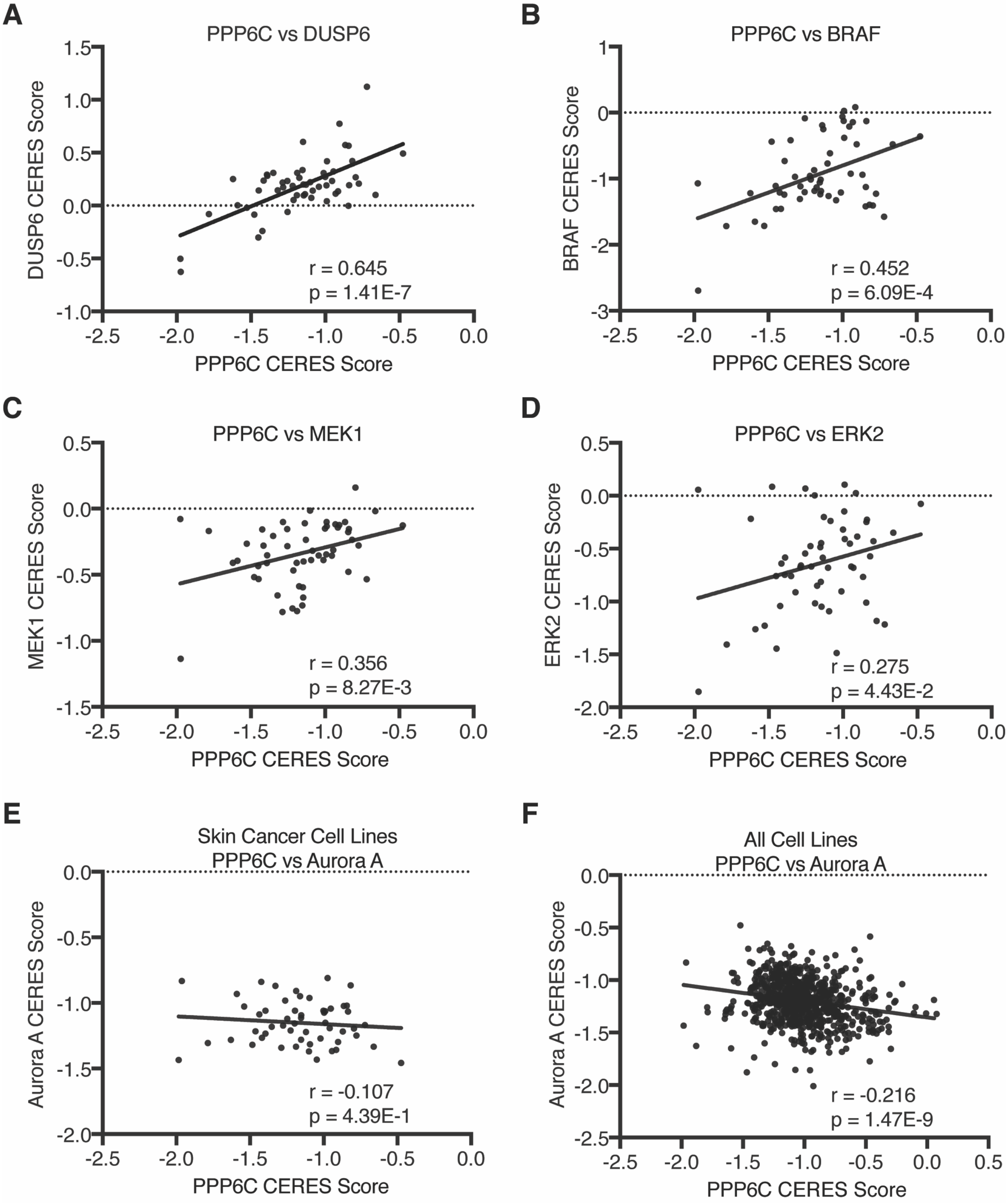


**Figure S3. Correlations of PPP6C dependency and ERK pathway dependency**
(A-E) CERES scores for PPP6C (x-axis) plotted against CERES scores for (A) DUSP6, (B) BRAF, (C) MEK1, (D) ERK2, and (E) Aurora A (y-axis). CERES scores are for all skin cancer cell lines from the Cancer Dependency MAP Project. Pearson's correlation coefficients (r) and associated *p*-values from linear regression analysis are indicated.

(A) CERES scores for PPP6C (x-axis) plotted against CERES scores for Aurora A (y-axis). CERES scores are for all cancer cell lines from the Cancer Dependency MAP Project. Pearson's correlation coefficient (r) and associated p-value from linear regression analysis are indicated.

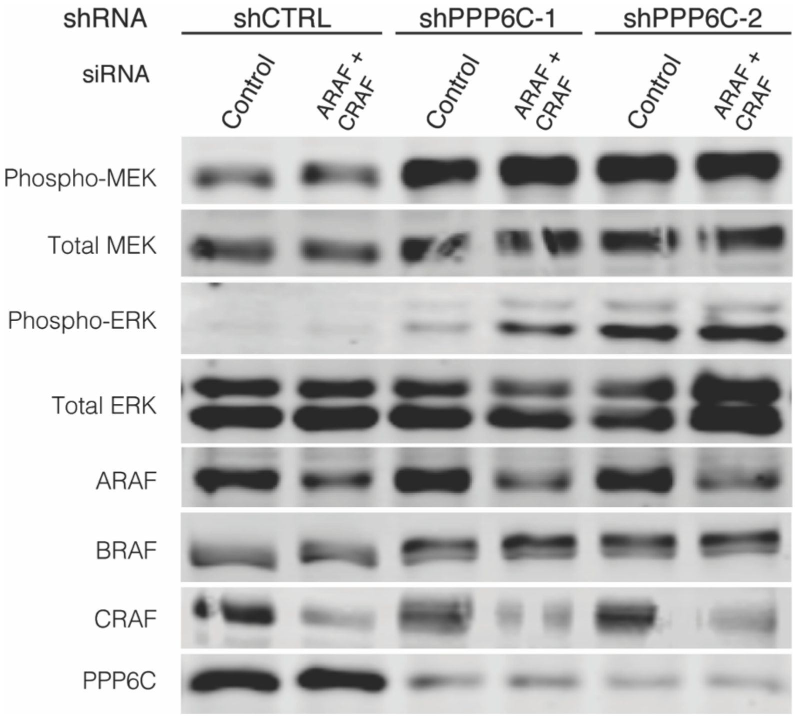


**Figure S4. ARAF and CRAF combined knockdown does not alter ERK hyperactivation by PPP6C loss**

(A) 501mel cells expressing shCTRL, shPPP6C-1, or shPPP6C-2 were transfected with non-targeting control siRNA or siRNA targeting both ARAF and CRAF. Cells were lysed and assessed by immunoblot for phosphorylated and total MEK and ERK. Knockdown of ARAF, CRAF, and PPP6C was also confirmed via immunoblot.

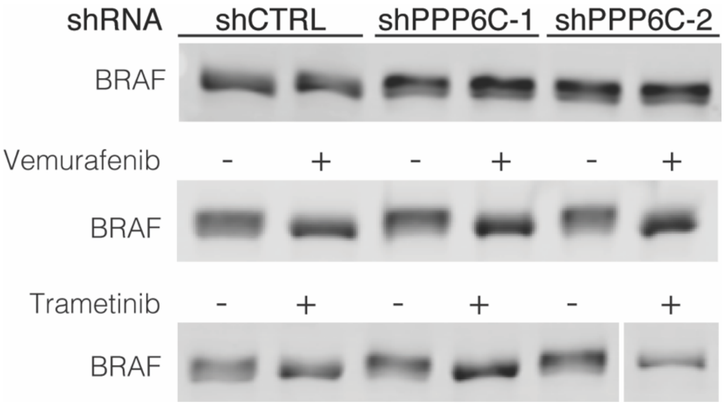


**Figure S5. PPP6C knockdown increases feedback phosphorylation of BRAF**
(A) shCTRL, shPPP6C-1, and shPPP6C-2 501mel cells were treated with 1uM vemurafenib or 50nM trametinib for 24 hours as indicated. Cells were lysed and assessed by immunoblot for BRAF electrophoretic mobility shifts indicative of changes in phosphorylation.

**Table S1. Raw Illumina sequencing read counts from shRNA screens**

**Table S2. RIGER analysis of shRNA screen results**
